# Connecting structure and function from organisms to molecules in small animal symbioses through chemo-histo-tomography

**DOI:** 10.1101/2020.09.28.316802

**Authors:** Benedikt Geier, Janina Oetjen, Bernhard Ruthensteiner, Maxim Polikarpov, Harald Gruber-Vodicka, Manuel Liebeke

**Author notes:** Corresponding Authors* Dr. Manuel Liebeke, Department of Symbiosis, Max Planck Institute for Marine Microbiology, Celsiusstraße 1, 28359 Bremen, Germany, Dr. Benedikt Geier, Department of Symbiosis, Max Planck Institute for Marine Microbiology, Celsiusstraße 1, 28359 Bremen, Germany. Notes* The authors declare no competing financial interest.

## Abstract

Our understanding of metabolic interactions between small symbiotic animals and bacteria or parasitic eukaryotes that reside within their body is extremely limited. This gap in knowledge originates from a methodological challenge, namely to connect histological changes in host tissues induced by beneficial and parasitic (micro)organisms to the underlying metabolites. To close this gap, we developed chemo-histo-tomography (CHEMHIST), a culture-independent approach to connect anatomic structure and metabolic function in millimeter-sized symbiotic animals. CHEMHIST combines spatial metabolomics based on mass spectrometry imaging (MSI) and microanatomy-based micro-computed X-ray tomography (microCT) on the same animal. Both high-resolution MSI and microCT allowed us to correlate the distribution of metabolites to the same animal’s three-dimensional (3D) histology down to sub-micrometer resolutions. Our protocol is compatible with tissue specific DNA sequencing and fluorescence in situ hybridization (FISH) for the taxonomic identification and localization of the associated micro(organisms). Building CHEMHIST upon in situ imaging, we sampled an earthworm from its natural habitat and created an interactive 3D model of its physical and chemical interactions with bacteria and parasitic nematodes in its tissues. Combining MSI and microCT, we introduce a workflow to connect metabolic and anatomic phenotypes of small symbiotic animals that often represent keystone species for ecosystem-functioning.

**Significance:** Metabolites mediate the establishment and persistence of most inter-kingdom symbioses. Still, to pinpoint the metabolites each partner displays upon interaction remains the biggest challenge in studying multi-organismal assemblages. Addressing this challenge, we developed a correlative imaging workflow to connect the in situ production of metabolites with the organ-scale and cellular 3D distributions of mutualistic and pathogenic (micro)organisms in the same host animal. Combining mass spectrometry imaging and micro-computed X-ray tomography provided a culture-independent approach, which is essential to include the full spectrum of naturally occurring interactions. To introduce the potential of combining high-resolution tomography with metabolite imaging, we resolve the metabolic interactions between an invertebrate host, its symbiotic bacteria and tissue parasites at unprecedented detail for model and non-model symbioses.

## Introduction

Earthworms represent a prime example of a keystone species (1) that experiences constant chemical interactions with bacteria (2), fungi (3), plants and small invertebrates (4) across soil ecosystems. Even within their tissues, earthworms harbor symbiotic microbes (5) and small animal parasites (6) that trigger internal metabolic responses such as innate immunity.

Unlike the metabolites involved in the earthworms’ digestion of leaf litter (7), the metabolites involved in the chemical interactions between earthworms and their associated (micro)organisms are unknown. For instance, most lumbricid earthworms harbor species-specific bacteria in their excretory organs. Still, it is unclear if the symbionts complement the host through vitamins or detoxification of nitrogenous waste products (8). In addition to mutualistic bacteria, nematodes infest the muscles, blood vessels and excretory organs in over ten earthworm species (9). The metabolic interactions between earthworms and their associated partners provide a versatile model to investigate how metabolites enable their role as engineers and janitors of soil ecosystems across the globe (4, 10).

The sum of mutualistic, commensal and pathogenic interactions results in a unique anatomic and in particular, metabolic phenotype for nearly every host individual (11, 12). Resolving this variability, insitu imaging of both metabolic and cellular phenotypes of the same host organ revealed metabolites which drive metabolic heterogeneity of the symbiotic partners (13, 14). For instance, correlative chemical imaging of the respiratory epithelia in a symbiotic invertebrate showed that within tens of micrometers the same species of intracellular bacterial symbionts produces different membrane lipids (13). Notably, metabolic interactions between animals and their microbes are not restricted to symbiotic tissues (15). Along the gut-brain axis, microbial metabolites produced in the gut can affect tissues across the host reaching the brain (16). Therefore, extending correlative chemical imaging into 3D approaches can be crucial for capturing how metabolites, involved in symbiotic interactions, distribute in host animals (17).

Previous studies have addressed this methodological challenge by combining non-destructive magnet resonance tomography (MRT) with matrix-assisted laser desorption/ionization (MALDI)-MSI, which enabled colocalizations between spatial chemistry and 3D anatomy of organs (18) and pathogenic abscesses (19-21). The spatial resolution used in these approaches was ideal for imaging animals with millimeter-sized organs such as mice. However, the majority of animals used as symbiosis models (22) aside medical studies have body sizes of only a few millimeters. Imaging their 3D anatomy and associated (micro)organisms together with their metabolite profiles requires each imaging technique to achieve micro- to nanometer resolutions. To capture the metabolic interactions that occur at the interface between host and associated (micro)organisms in situ imaging of the symbiotic tissues is essential.

The integration of micro-computed tomography (microCT) and MALDI-MSI technologies is an emerging approach to image an animal’s 3D histology and spatial chemistry at micrometer scales (23). MicroCT is a non-invasive approach allowing X-ray imaging of 3D histology, and unlike MRT, microCT can reach subcellular resolution (24-27). For metabolite imaging, MALDI-MSI techniques have also reached subcellular resolutions (28, 29). For imaging animal models, both techniques have been mainly applied independently based on their different imaging workflows. The principal obstacles are that MSI requires tissue sectioning and thus cannot be applied before non-destructive 3D imaging. Conventional microCT on the other hand requires chemical contrasting of soft tissues (30), which would change the chemistry of the sample and interfere with subsequent MSI (31).

Here we present **chem**o-**his**to-**t**omography (CHEMHIST) that combines MALDI-MSI and microCT to image both the spatial chemistry and 3D microanatomy of the same small symbiotic animal. Our objective was to make CHEMHIST also applicable to animals directly sampled from their natural habitat through state-of-the-art in situ imaging and metagenomic DNA sequencing within one pipeline. CHEMHIST provides an up to two orders of magnitude higher resolution than previous correlative 3D MALDI-MSI approaches. This advance allowed us to take an earthworm from the environment and create a 3D atlas of its chemical and physical interactions with bacteria and nematodes naturally occurring inside its tissues.

## Results and Discussion

We used earthworms as target organisms for developing a correlative high-resolution tomography and metabolite imaging workflow. They are easily accessible and their diverse associations with parasites and microbes result in phenotypic heterogeneity that demands correlative imaging at scales from millimeters to micrometers. Their anatomical segmentation presents a modularly repeated anatomy that facilitates the identification of irregularities in the chemical and anatomic phenotype of the animal.

### CHEMHIST workflow for creating a multimodal 3D atlas

CHEMHIST consists of three major steps. First, we physically divided the sample for the different fixation and sample preparation procedures of MSI and microCT (Fig. 1A). Second, we created a combined 3D overview of the spatial chemistry and anatomy of the animal tissue. Applying MSI and microCT at moderate instead of high-spatial resolutions was sufficient to resolve organs from tens to hundreds of micrometers and allowed faster imaging at larger fields of view (e.g. whole cross sections of an animal with MSI). We combined the imaging data to create a 3D model of the spatial chemistry and anatomy at a whole animal scale. In the third step, we analyzed the combined chemo-histo 3D model to determine regions of interest focusing on tissues that were colonized by microbes or parasites to guide subsequent high-resolution MSI and microCT. Because we used the CHEMHIST 3D model as an overview to guide the high-resolution measurements, we referred to the 3D model as an atlas (32).

**Fig 1:**
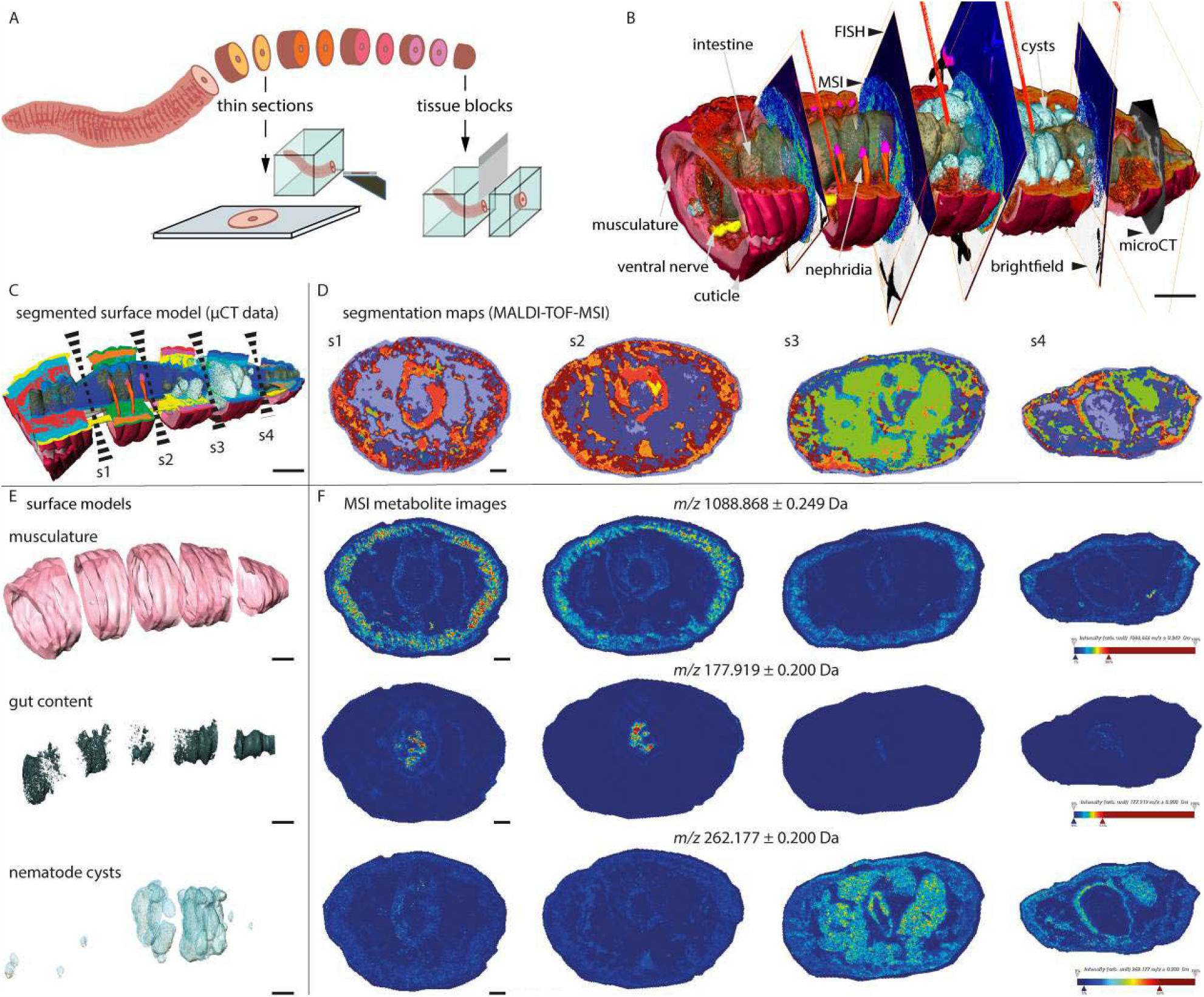
Chemo-histo-tomography (CHEMHIST) revealed organ specific chemistry in the posterior segments of an earthworm. **A)** Division of the animal into alternating tissue sections for spatial metabolomics and microscopy and tissue blocks for tomography. **B)** 3D CHEMHIST atlas at organ-scale with data from microCT, MALDI-TOF-MSI, FISH and brightfield microscopy. **C)** Segmentation of the 3D microCT data (surface models, semi-automatic segmentation) and **D)** 2D MALDI-TOF-MSI data to delineate the spatial chemistry (unsupervised spatial metabolite clustering). Sectioning planes of sections s1–s4 indicated in **C** with dashed wedge. **E)** Examples of surface models of individual organs and **F**) individual metabolites located in the organ throughout sections s1–s4. Scale bars in **B, C** and **E**: 2000 µm (2D scale bars in 3D models as approximate scale) and magnifications in **F**: 500 µm.

To apply MSI and microCT to the same animal, here an earthworm (*Lumbricus rubellus*), we had to treat the sample in a manner that preserves both morphology and spatial chemistry, but without one technique interfering with the other. For CHEMHIST, snap freezing provided a tradeoff between preserving enough anatomic details for microCT without modifying the spatial chemistry for MALDI-MSI. For microCT and MALDI-MSI we divided our frozen sample into two sample types: tissue blocks and tissue sections. The tissue blocks of one to three millimeter thickness were trimmed off the frozen sample with a razor blade and tissue sections of 16 µm thickness were sectioned off each tissue block with a cryotome (Fig. 1A). Choosing cross sections over longitudinal sections allowed us to increase the z-resolution and make our sectioning protocol applicable to the majority of bilateral symmetrically organisms. We obtained tissue blocks for microCT and tissue sections for MSI. Additionally, we stored consecutive tissue sections from in between the tissue blocks for high-resolution MSI and spatially targeted metagenomics sequencing. Obtaining both sample types in an alternating manner provided sample pairs of one tissue block for microCT and one adjacent tissue section for MSI (Fig. 1A). Sharing the same sectioning interface, each tissue section anatomically and chemically matched its adjacent tissue block, which allowed us to precisely correlate microCT and MALDI-MSI data despite segregating the tissues.

To generate an anatomic 3D atlas, we imaged each of the five tissue blocks with microCT at a 4.4 µm voxel size. From the five microCT datasets, we virtually reconstructed a 3D anatomy model of the specimen (Fig. 1B and 3A, *SI Appendix*, Fig. S-2). To gain an overview of the spatial chemistry throughout the sample, we imaged four cryo-sections with MALDI time-of-flight (TOF)-MSI at a 25-µm pixel size (*SI Appendix*, Fig. S-3). After MALDI-TOF-MSI we used brightfield microscopy and fluorescence in situ hybridization (FISH) to image the histology and the bacteria in each of the four tissue sections (13)(Fig. 1B and *SI Appendix*, Fig. S-4). To complete the 3D CHEMHIST atlas, we co-registered the 2D imaging MALDI-TOF-MSI, FISH and brightfield microscopy datasets into the anatomic 3D model (Fig. 1B, *Appendix*, Fig. S-2). The 3D imaging platform AMIRA© provided a graphical user interface (GUI) to visualize and co-register 3D and 2D imaging data (*SI Appendix*, Video S1). Using GUI-based software allows non-computer scientists to integrate their own correlative imaging data without programming and readily enables 3D data exploration and analysis.

### CHEMHIST provides a cross-kingdom link between spatial chemistry and 3D anatomy

To study the association between organ-specific metabolites and 3D anatomy of the earthworm, we needed to transform the data of the different techniques into a comparable format. Unsupervised spatial clustering of the metabolite images (33) and semi-automatic thresholding of anatomic features in the 3D microCT data resulted in binary segmentation maps of both data types (Fig. 1C–F). Delineating both the MSI and microCT data through different segmentation strategies provided a comparison of anatomic 3D structure and metabolic function.

Organ-specific anatomy as well as chemistry are inherently linked to organ functioning, such as in movement, digestion or signal transduction. In the dataset of the earthworm, we consistently located specific metabolites in each tissue cross-section (Fig. 1F). For example, in the musculature, we located lombricine with (AP)-MALDI-orbitrap-MSI (*m/z* 271.0802, [C_6_H_15_N_4_O_6_P□+□H]□) an annelid specific energy storage metabolite (34) (*SI Appendix*, Table S-1). In the earthworm’s body wall, we detected protoporphyrin (*m/z* 563.2653, [C_34_H_34_N_4_O_4_□+□H]□), a typical pigment in the dorsal epidermis of lumbricid earthworms (35) (*SI Appendix*, Table S-1). Detecting metabolites in continuous organs over the length of the animal, such as musculature and gut content could reveal metabolic heterogeneity indicating changes in the microenvironments along the body axis of the animal (Fig. 1E).

The key advantage of combining microCT with MSI was to image tissue volumes at a resolution sufficient to identify discontinuous organs and anatomic abnormalities and target them with MSI. For instance, the nephridia, which occur pairwise in each segment and harbor symbiotic bacteria (5, 8), were not present in each tissue section used for MSI (Fig. 2). The 3D CHEMHIST atlas helped to locate histological structures that originated from sectioning planes through the nephridia (Fig. 1B and 2A). Applying correlative FISH microscopy on the same tissue sections after MSI (13) with probes targeting eubacteria allowed us to locate bacterial accumulations across each tissue section and identify the symbiotic tissues of the nephridia (Fig. 2A, D and E). We used the fluorescence signals of the bacteria in the same tissue section after MSI to screen for metabolites that spatially correlated to the bacterial accumulations in the nephridia identified from the 3D atlas (Fig. 2B and C) (*SI Appendix*, Fig. S-4) (13).

**Fig 2:**
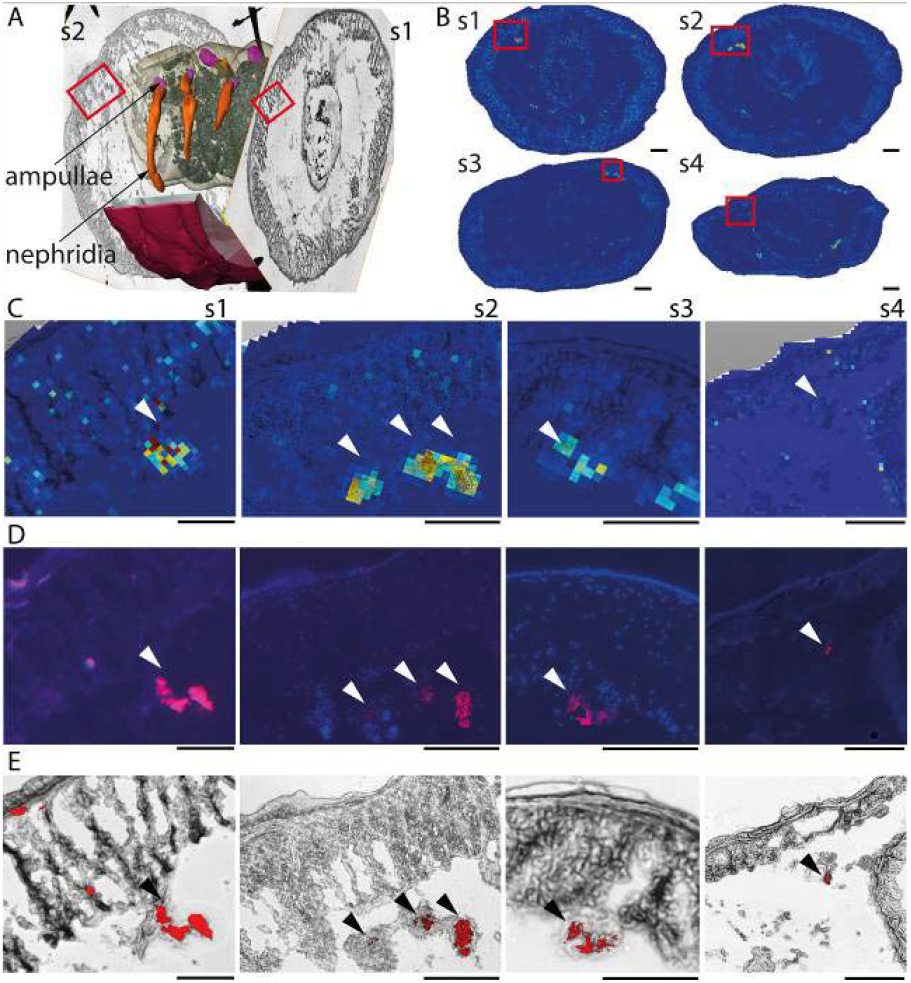
Bacterial cells within organs have unique molecular fingerprints. **A)** The 3D model shows the surface reconstruction of the nephridia, including the bacteria-containing ampullae of the second tissue piece between tissue section s1 and s2. **B)** The ion images of whole tissue sections show the metabolite *m/z* 1116.833, found through the colocalization analysis of MALDI-MSI and FISH signals. Red boxes in s1–s4 indicate magnified areas, shown as MSI and bright field overlay in **C). D)** FISH microscopy images and **E)** FISH and brightfield overlay with bacteria labeled in red. Scale bars of whole tissue sections s1–s4 (**B**): 500 µm and magnifications in **C–E**: 250 µm.

### Guiding high-resolution MSI and microCT to visualize host–parasite interactions with CHEMHIST

The 3D CHEMHIST atlas of a field-collected earthworm also allowed us to study structures that are not part of an earthworm’s anatomic bauplan (blueprint) in detail. Our analysis of the 3D atlas indicated 20-30 µm (in diameter) sized parasitic worms in the earthworm tissues. We chose these small worms to showcase how CHEMHIST can be used to integrate micrometer-scale metabolite imaging, nanometer-scale histology and metagenomics sequencing to resolve the metabolic and anatomic phenotypes of these parasites in situ (Fig. 3).

**Fig 3:**
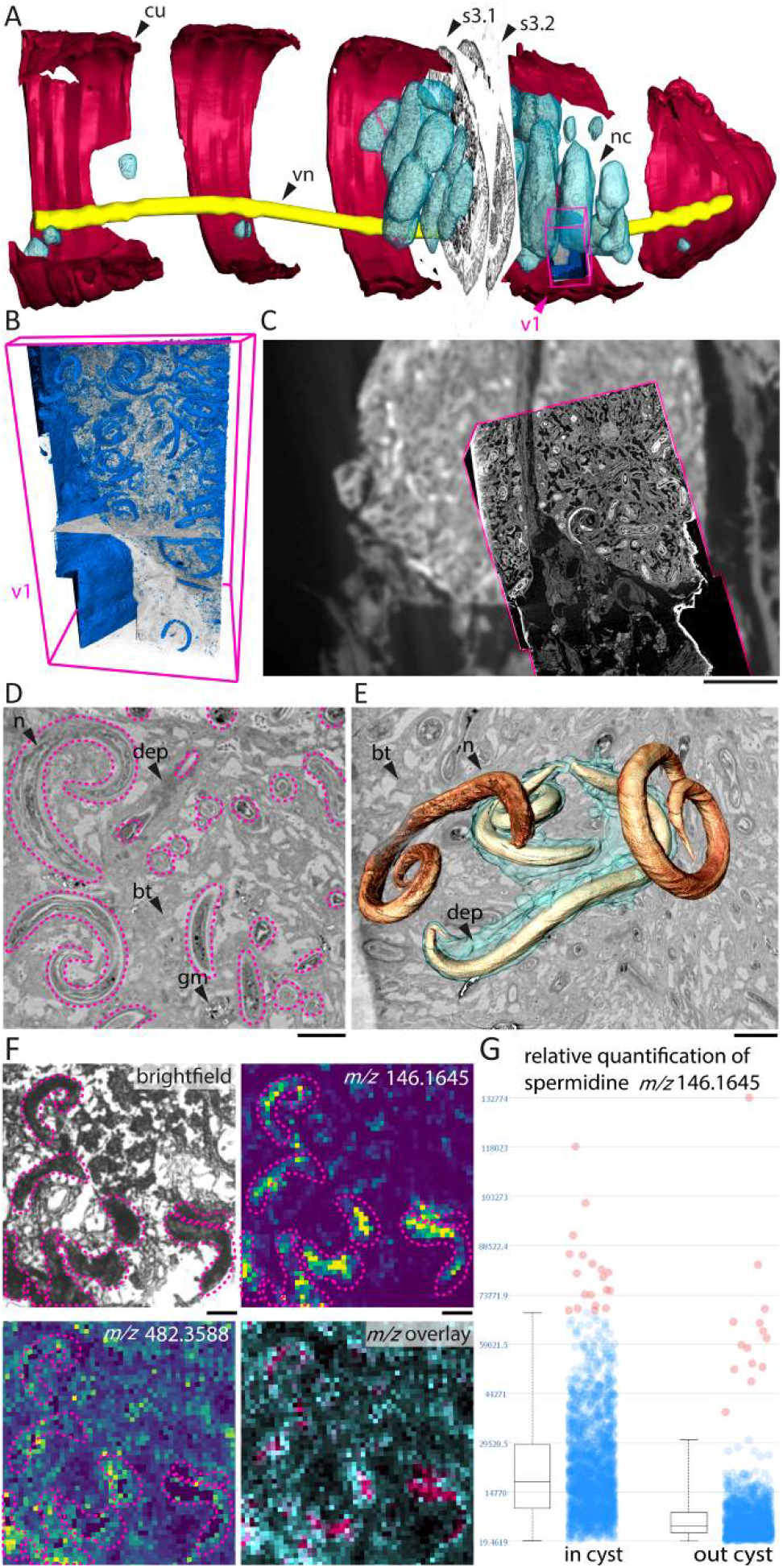
Using the 3D CHEMHIST atlas to guide high-resolution imaging of the interactions between earthworm and parasitic nematodes. **A)** MicroCT model with the co-registered MSI sections (s3.1 and s3.2) and the tissue volume (v1) imaged with high-resolution SRmicroCT. **B)** Isosurface 3D rendering of the nematodes (blue) and a virtual sectioning plane (*xy*) through the SRmicroCT image stack (v1). **C)** Overlay of the microCT and SRmicroCT (magenta outline, *xy*-plane) to show the increased resolution and detail gained with SRmicroCT. **D)** Virtual plane through the SRmicroCT data shows sections of nematodes in the cysts (magenta outlines) **E)** 3D renderings of four nematodes of which two were surrounded by a homogeneous deposit (cyan cloud). **F)** high resolution MSI shows distribution of two metabolites, for orientation the bright field image of section s3.2 shows magnified the nematodes (magenta outlines) in a cyst. The distributions of spermidine (*m/z* 146.1645) and PAF (*m/z* 482.3588) and an overlay of both metabolites is shown (PAF in cyan, spermidine in magenta). **G)** Relative quantification of spermidine colocalized with nematode tissues (outlined in F and *SI Appendix*, Fig. S-8) in- and outside the brown body cysts. **cu**, cuticle; **nc**, nematode cyst; **vn**, ventral nerve cord; **n**, nematode; **dep**, deposit; **bt**, brown body tissue; **gm**, granular mass; **i**, electron dense inclusions. Scale bars in C: 500 µm, in D, E and F: 50 µm.

In the 3D atlas the cysts containing the nematodes occurred irregularly, increasing in size and density towards the posterior of the earthworm (Fig. 1B and 3A, *SI Appendix*, Fig. S-6). By specifically targeting cyst tissues for DNA sequencing and phylogenetic analysis, we identified the nematodes as *Rhabditis maupasi* (*SI Appendix*, Fig. S-5). The species resides as parasites in the nephridia and coelomic cavities in common earthworm species such as *Lumbricus terrestris, Lumbricus rubellus, Allobophora longa*, and *Allobophora turgid* (36).

In earthworms, these cysts are called brown bodies and are produced to encapsulate and degrade organic debris, microbes and parasites through reactive oxygen species (37, 38). To visualize the nanometer-scale 3D histology during the degradation process of single nematodes in the brown bodies, we used synchrotron radiation-based (SR)microCT. This high-resolution microCT technique allowed us to rescan cysts areas with high densities of nematodes at an eleven times increased resolution (0.325 µm voxel size) (Fig. 3A and B).

In the brown body tissue, SRmicroCT we could visualize distinct histopathological states of the nematodes, only known from stereomicroscopic observations of live animals (39). We found nematodes that contained highly electron-dense inclusions and nematodes that were in the process of disintegrating into a granular mass, possibly induced by the earthworm’s immune response (Fig. 3D and E, *SI Appendix*, Fig. S-6) (39). We also found histologically intact nematodes, some of which were surrounded by a homogeneous deposit (Fig. 3D and E). Based on electron microscopy, this deposit was hypothesized to be a humoral response of the earthworm against the nematodes in the coelomic cavity (6).

Nearly five decades later, our integrated micrometer-scale metabolite imaging also provides insights into the potentially metabolic function of the earthworm’s humoral response. The nematode-filled cysts displayed a unique metabolome, characterized by a specific metabolite cluster originating from the host-parasite interaction (Fig. 1B, C, F and 2A). To guide metabolite imaging of single nematodes, we chose a consecutive tissue section to the MALDI-TOF MSI section with nematode cysts (Fig. 3F and *SI Appendix*, Fig. S-7). We imaged the encysted nematodes with atmospheric pressure (AP)-MALDI-orbitrap-MSI, a high-resolution MSI technique that provided a three-time higher spatial resolution of 8-µm pixel size. The high mass accuracy of the orbitrap detector (below 3 ppm), allowed for more precise annotations of the detected metabolites. For data analysis and exploration, we co-registered both high-resolution datasets, SRmicroCT and AP-MALDI-orbitrap-MSI, into the 3D atlas extending the 3D model with different levels of resolution of the same structures (Fig. 3A).

The metabolite images revealed that most nematodes were surrounded by a platelet activation factor (PAF), specifically lysophosphatidylcholine (lysoPC)O-16:0/0:0 (*m/z* 482.3605, [C_24_H_52_NO_6_P□+□H]□) (Fig. 3F, *SI Appendix*, Fig. S-7 and Table S-1). PAFs are single fatty acid chain phospholipids that serve as inflammatory modulators, conserved in metazoans across all domains of life (40). However, beyond the up- and downregulation of PAF lipids as humoral immune response, their site of production upon animal–microbe and animal–parasite interactions remains unknown (41, 42). The lysoPC(O-16:0/0:0) that we detected accumulating around the nematodes likely promotes the aggregation of hemocytes (43), which form most of the brown body tissue (6, 38) and release the reactive oxygen species (42). Although we hypothesize that the earthworm produces PAF lipids as an inflammatory response to the nematodes, solely based on our imaging data, we cannot exclude that the PAFs originated form the nematodes. Nevertheless, we show that PAF lipids concentrate at the host–parasite interface, whereas they are absent in the nematode tissues, but colocalize with aggregated hemocytes (*SI Appendix*, Fig. S-7).

Focusing on the metabolite profiles of encysted nematodes, we detected twice as much of the polyamine spermidine (*m/z* 146.1652, [C_7_H_19_N_3_□+□H]□) compared to nematodes not encapsulated within the brown bodies. Not encysted nematodes had spermidine signals as low as earthworm tissues (Fig. 3G, *SI Appendix*, Fig. S-7, S-8 and Table S-1). Spermidine supplementation in other nematodes enhances longevity, inducing autophagy and suppressing oxidative stress and necrosis (44). The nematodes encapsulated in brown bodies might increase their levels of spermidine to inhibit necrosis, as a protection against the reactive oxygen species of the earthworm. Nematodes within brown bodies showed.. The production of spermidine as an anti-oxidative stress response could help some of the nematodes to survive in the brown bodies until the earthworm sheds its posterior segments (45) providing an escape mechanism from their host (39, 46).

## Conclusion

Faced with the intimidating complexity of natural systems, scientists have studied model organisms under controlled conditions and so gained an understanding of their detailed molecular biology. This meticulous research on model organisms has created a strong foundation of databases and new technologies. Today, this groundwork allows us to leave the laboratory and tease apart the array of metabolic interactions between organisms and their effects on phenotypic heterogeneity in nature. CHEMHIST offers an unprecedented in-depth visualization of such interactions between millimeter-sized animals and their (micro)organisms in situ and without prior knowledge of the sample. Notably, CHEMHIST requires an elaborate workflow and access to different devices and it is thus challenging for larger sample sizes (e.g. hundreds of specimens). The strength of CHEMHIST is to enable a comprehensive metabolomics data exploration of interacting organisms on a cellular level.

Extending histotomography (26) into chemo-histo-tomography (CHEMHIST) opens nearly uncharted territory for label-free correlative imaging. Although our approach provided a wealth of biological and biochemical information, our approach of dividing the sample for the separate techniques led to a minimal loss of tissue and potentially relevant details. Future development of CHEMHIST could integrate phase-contrast SRmicroCT (47), an emerging technique that allows quantitative 3D imaging of tissues without contrasting agents at nanometer scales (48, 49). Serial MSI after phase-contrast SRmicroCT would provide data for a lossless anatomic and metabolic 3D model of the same organism (50) and the basis for automated correlations between modalities in 3D (51).

We envision that our advances in correlative chemical and structural in situ imaging will drive discovery-based research and fuel scientists’ hypotheses on the metabolic interactions of their symbiotic systems.

## Materials and Methods

### Chemicals and reagents

All chemicals were obtained from Sigma-Aldrich (Steinheim, Germany) unless specified otherwise.

### Tissues and sample collection

An adult earthworm (*Lumbricus rubellus*) was collected from soil in a polder region of the central Netherlands (supplied by Lasebo BV, Netherlands) and rapidly frozen in isopentane cooled with liquid nitrogen. The posterior end (30 segments, approx. 2 cm) was embedded in 2% carboxymethylcellulose (CMC) gel and subsequently solidified at −20 °C. Using a precooled razor blade, the embedded earthworm was trimmed inside a cryo-chamber. Sectioning of the frozen sample block was performed as follows. The first tissue section (1–3 mm) was cut with a razor blade from the frozen CMC block. The block was then trimmed and several 16 µm sections were obtained with a cryotome (−20 °C). This was repeated to obtain five blocks of tissue and four adjacent thin sections. Thin sections were transferred onto Bruker® ITO glass slides via thaw mounting. Small crosses (1–2 mm) were drawn around the dried samples using a white paint marker (edding ®751) as a reference for the computational alignment (14). The slides were stored at 4 °C prior to MSI matrix application. The tissue blocks were used for microCT measurements and the adjacent thin sections for correlative brightfield microscopy, MSI and FISH imaging.

### Micro-computed tomography

The frozen tissue blocks were defrosted in 8% paraformaldehyde. This chemical post-cryo fixation with paraformaldehyde and osmium tetroxide allowed us to minimize the tissue damage induced by thawing and preserve subcellular detail for high-resolution microCT. Contrasting was achieved using an aqueous 1% osmium tetroxide/acetone (1:1 vol./vol.) solution for 2.5 h at 20 °C. Dehydration and infiltration were conducted according to the 45345 FLUKA Epoxy embedding medium data sheet. The resin blocks were pre-trimmed with a fretsaw, then fine trimmed using a razor blade (52). Trimmed embedded earthworm tissue blocks were mounted individually on glass rods with a hot-melt gun. A “nanotom m” computed tomography system (GE Measurement & Control, Wunstorf, Germany) was used for the microCT analysis of the tissue blocks with the following X-ray parameters: 110 kV, 120 μA, 0.75 s exposure time, averaging = 4, scanning time = 1.5 h, number of projections acquired during scan = 1500. The tomographic reconstruction was performed using the phoenix datos|x 2.2 reconstruction software (GE Measurement & Control, Wunstorf, Germany) on a separate workstation and resulted in a voxel size of 4.4 µm. The 16-bit volume is saved in *.vgl format. This format was imported into the 3D-visualization software VGStudio (2.2) (Volume Graphics, Heidelberg). Cropping, histogram adjustments, bit dept (to 8 bit) and format (to *.raw volume) conversions were performed in VGStudio.

### Synchrotron radiation-based micro-computed phase-contrast X-ray tomography P14 beamline

After determining the position of the encysted nematodes in the laboratory-based microCT data, one small tissue block (∼ 1 × 1 × 3 mm) was cropped out from tissue block 3 with a razor blade in an area that contained the nematodes. The cropped out block was mounted onto one SPINE sample holder (53). Experiments were carried out on the EMBL undulator beamline P14 at the PETRA-III storage ring (c/o DESY, Hamburg, Germany) using the propagation-based phase-contrast imaging setup described in (54, 55). X-ray energy of 18 keV was used. The X-ray images were obtained using an X-ray microscope (Optique Peter, Lyon, France) consisting of an LSO:Tb scintillator with a thin active layer of 8 µm, an OLYMPUS UPlanFL 20-fold objective lens (Olympus, Tokyo, Japan), a 45°-reflecting mirror, an OLYMPUS 180 mm Tube Lens and PCO.edge 4.2 sCMOS camera with a 2048 × 2048 pixels sensor at a pixel size of 6.5 µm. The effective pixel size of 0.325 µm resulted in a 666 × 666 µm^2^ field of view. To ensure artifact-less phase retrieval (56) in the near-field edge-enhancing regime, each tomographic acquisition consisted of four measurements at sample-to-detector distances of 5.9, 6.4, 7, and 7.9 cm. A total of 1850 projections covering 185 degrees of continuous rotation and 40 flat-field images were acquired at each distance with a frame rate of 100 frames per second. A complete four-distance tomographic data acquisition took less than 2 minutes. Three tomograms with overlapping areas were acquired along the vertically shifted sample.

### SRmicroCT Data processing

Data processing was carried out using in-house Python software, performing flat-field correction, phase retrieval and tomographic reconstruction. First, each X-ray image of the sample was divided by the flat-field image with the highest similarity. For this operation, we used the similarity index (SSIM) implemented in the scikit-image Python module as a metric (57). Subsequently, a four-distance non-iterative holographic reconstruction procedure (58) was applied with a δ/β ratio of 0.17 to obtain a projected phase map of the sample at the given angle. The tomographic reconstruction was then performed using the tomopy Python module (59) with ‘gridrec’ algorithm and ‘shepp-logan’ filter. The three reconstructed volumes were co-registered with the commercial software Amira 6.7.0 (Thermo Fisher Scientific, USA), which did not require any non-linear operations, because we used the transform editor.

### MALDI-TOF-MSI

For MALDI-MSI, a matrix consisting of 7 mg mL^-1^ α-cyano-4-hydroxycinnamic acid in 70:30 acetonitrile/water with 0.2% trifluoroacetic acid was applied via an automated spray-coating system (SunCollect, SunChrom GmbH, Germany) using the following parameters: *z*-distance of the capillary 25 mm; pressure of compressed air 2 bar; the flow for the first layer was 15 µL min^-1^ and for layers 2– 8 20 µl min^-1^. MALDI-MS imaging was performed using an Autoflex speed™ LRF MALDI-TOF (Bruker Daltonik, Germany) with MALDI Perpetual™ ion source and smartbeam™-II 1 kHz laser and reflector analysis in positive modes (18). A spot size of 25 µm was used and 500 shots per sampling point acquired using a “random walk” pattern with 100 shots per location within the sampling spot. The mass detection range was set to *m/z* 100–1280 with 200 ppm accuracy. For data processing and visualization of the MALDI-MS imaging data, flexImaging™ 4.0 (Bruker Daltonik, Germany, 2015) was used.

### AP-MALDI-orbitrap-MSI

High spatial resolution MSI was performed using an atmospheric-pressure scanning microprobe MALDI source (AP-SMALDI10®, TransMIT GmbH, Giessen, Germany) with a Q Exactive™ plus Fourier transform orbital trapping mass spectrometer (Thermo Scientific, Germany). A nitrogen laser with a wavelength of 337 nm and 60 Hz repletion rate was used for desorption and ionization. The laser beam was focused to an 8 µm ablation spot diameter and a step size of 8 µm in *x* and *y* was used to scan the sample. Mass spectral acquisition was performed in positive mode and *m/z* 100–1000 Da with a mass resolving power of 140,000 (dataset in Fig. 3) at 200 *m/z* with a mass accuracy <5 ppm. Additional datasets for the annotation of metabolites in METASPACE (60) and MS/MS experiments were recorded with a resolving power of 240,000 (see *SI Appendix* Table S-1) at 200 *m/z* with a mass accuracy <5 ppm. The mass spectrometer was set to automatic gain control, fixed to 500 ms injection time. For data processing and visualization, ImageQuest 1.1 (Thermo Scientific) was used.

### MALDI-MS^2^

The identification of spermidine and PAF was supported by MALDI-MS^2^ experiments (*SI Appendix*, Table S-1 and Figure S-9). For PAF we obtained enough ions to use on tissue MALDI-MS^2^ in positive-ion mode with the mass analyzer set to a resolution of 240k resolution and a collision energy of 25 eV in the HCD cell. For spermidine, we could not obtain sufficient ions from the tissue to perform on tissue fragmentation. To support the identification of the exact mass, we matched the MS^1^ *m/z* values of spermidine measured from the tissue with the MS^1^ *m/z* values from a spermidine standard. The standard was spotted onto a glass slide and fragmented with MALDI-MS^2^, in positive-ion mode with the mass analyzer set to a resolution of 240.000 resolution and a collision energy of 30 eV in the HCD cell.

### MALDI-MSI data analysis and visualization

The *.raw files were centroided and converted to *.mzML with MSConvert GUI (ProteoWizard, version 3.0.9810 (61)) and then to *.imzML using the imzML Converter 1.3 (62). SCiLS Lab software (SCiLS, Bruker Daltonik GmbH, Bremen, Germany) version 2019b was used for spatial segmentation analysis and alignment of brightfield microscopy and MSI datasets as a template for ROI selection.

### DNA extraction and metagenomics sequencing

Genomic DNA was extracted using the DNeasy Blood & Tissue Kit (Qiagen, Hilden, Germany). In brief, the nematode cysts from one consecutive tissue section of tissue section s3 (*SI Appendix*, Fig. S-5) were scraped off the glass slide using a sterile scalpel and transferred into a tube containing 180 μl buffer ATL and 20 μl proteinase K. The tissue was digested at 56°C for 3 days. Subsequent extraction steps were performed according to the manufacturer’s instructions. In the end 100 μl elution buffer were applied to the column and incubated at room temperature for 10 min. After the first round of elution, a second elution 100 μl of buffer was performed and the two elutions were pooled. The extracted DNA was stored at 4°C until further processing.

Illumina-library preparation and sequencing were performed by the Max Planck Genome Centre. In brief, DNA quality was assessed with the Agilent 2100 Bioanalyzer (Agilent) and genomic DNA was fragmented to an average fragment size of 400□base pairs (bp). An Illumina-compatible library was prepared using the TPase-based DNA library protocol. One nanogram of genomic DNA was cut and specific sequences were introduced by the Illumina Tagment DNA Enzyme (Illumina). Products were amplified by KAPA 2G Robust polymerase (Roche) with 15 cycles to enrich and to add library-specific barcoding information to the PCR products. After quality check by LabChip GX II (Perkin Elmer) libraries were pooled and sequenced on an Illumina HiSeq3000 sequencer with 2 x 150 bp paired end mode. Three million 150□bp paired-end reads were sequenced on a HiSeq 3000 (Illumina).

### Parasitic nematode phylogenetic analyses using the small subunit rRNA gene

We used phyloFlash v3.3 beta1 (https://github.com/HRGV/phyloFlash) (63) to assemble full-length SSU genes from the metagenomic reads. The nematode SSU matrix was constructed from the assembled *Rhabditis* related sequence, the available full length SSU genes of all species level representatives of the *Rhabditis* group and *Teratorhabditis* sequences as an outgroup. The sequences were aligned using MAFFT v7.394 (64) in G-Insi mode. The phylogenetic tree was reconstructed using FastTree v2.1.5 (65) with a GTR model, 20 rate categories and Gamma20 likelihood optimization, generating approximate likelihood-ratio-test values for node support. The tree was drawn with Geneious R11 (https://www.geneious.com) and rooted with *Teratorhabditis* as an outgroup.

### Fluorescence in situ hybridization and brightfield microscopy

After MALDI-MS imaging, the matrix was removed by dipping the sample slide into 70% ethanol and 30% water (vol./vol.) for 1 min each. In a second step, the tissue sections were post-fixed for one hour at 4°C in 2% paraformaldehyde in PBS. The sample was dried under ambient conditions and then prepared for catalyzed reporter deposition fluorescence in situ hybridization (CARD-FISH) following ref (66). For in situ hybridization, general probes were used that target conserved regions of the 16S rRNA in bacteria (EUB338; I-III; I: 5’-GCT GCC TCC CGT AGG AGT-3’, II: 5’-GCA GCC ACC CGT AGG TGT-3’, III: 5’-GCT GCC ACC CGT AGG TGT-3’) (67, 68). To visualize tissue containing DNA, samples were stained for nuclei with DAPI for 10 min. at room temperature, washed three times for 1 minute and mounted in a VECTASHIELD®/Citiflour® mixture (2:11) with 1 part PBS (pH 9) under a cover slip (Menzel glass, 24 × 60 mm, #1.5). Image acquisition for individual photomicrographs was carried out using a Zeiss Axioplan 2 microscope.

To image large sections for brightfield and fluorescence microscopy at high resolution, an automated microscope (Zeiss Axio Imager Z2.m, 10× objective) with a tile-scan Macro for Axio Vision was used. Individual images were acquired with 15% overlap and stitched (Fiji Plugin stitching 1.1).

### Composition of the 3D atlas

To combine the different modalities into the 3D atlas, tools in Amira 6.7.0 were used for segmentation, surface rendering and 3D co-registration (52). The individual microCT volumes were imported as *.raw files and manually realigned, based on the estimations of the location and composition of shared morphological features. Exact spacing of the intervals between tissue blocks relied on an estimate, as the precise thickness of tissue used for the sections could not be recorded due to loss of tissue during sectioning. However, information about the overall number of the earthworm segments allowed a realistic reconstruction of the overall spatial relations of the specimen.

Subsequently, the light microscopy images were co-registered into the spaces between tissue blocks, using morphologic structures in the orthographic cross-section of the microCT data as a template. Based on the previous alignment of the 2D-imaging modalities in SCiLS Lab, the co-registration parameters could be applied to the FISH- and MALDI-imaging data. All 3D visualizations and applications with high computation demands were performed using a 3D-imaging workstation (Windows 7 Professional, 64 bit, Intel Core i7-5960X CPU with 16 processors × 3.5 GHz, 64 GB RAM and NVIDIA Quadro P6000 with 24 GB).

## Author Contributions

BG and ML conceived the study. BG prepared all the samples and together with BR acquired the laboratory based microCT. MP acquired and reconstructed the SRmicroCT data. 3D and 2D visualizations and the co-registrations in Amira were done by BG. HGV coordinated metagenomics sequencing and analyzed the data. JO acquired the MALDI-TOF-MSI datasets and BG acquired the AP-MALDI-orbitrap-MSI dataset and FISH microscopy. The analysis of the MSI data in SCiLS Lab was done by BG. The manuscript and the figures were drafted and edited by BG and ML. All authors edited and commented on the manuscript.

## Data availability

All datasets (MSI, microCT and microscopy) can be accessed and directly downloaded under: https://figshare.com/projects/CHEMHIST_connecting_structure_and_function_from_organisms_to_molecules_in_small_animal_symbioses_through_chemo-histo-tomography/73527

A video animation of the multimodal 3D atlas can be found here: https://figshare.com/articles/media/final_mpg/4003944

## Acknowledgements

We thank Maggie E. Sogin (MPI Bremen) for constructive feedback and Janine Beckmann (MPI Bremen) for help with MALDI-MSI and MS/MS experiments and Miriam Sadowski (MPI Bremen) for help with DNA extractions. We thank Russell Naisbit, Wolfgang Geier and Grace D’Angelo for additional edits of the manuscript draft. We thank Nicole Dubilier for giving access to resources and for constructive feedback. This work was funded by the Gordon and Betty Moore Foundation Marine Microbiology Initiative Investigator Award (Grant GBMF3811) and the Max Planck Society.

