## Supplementary material for "Connecting structure and function from organisms to molecules in small animal symbioses through chemo-histo-tomography"

*Corresponding author:

**This PDF file includes:**

Figures S1–S22

Table S1

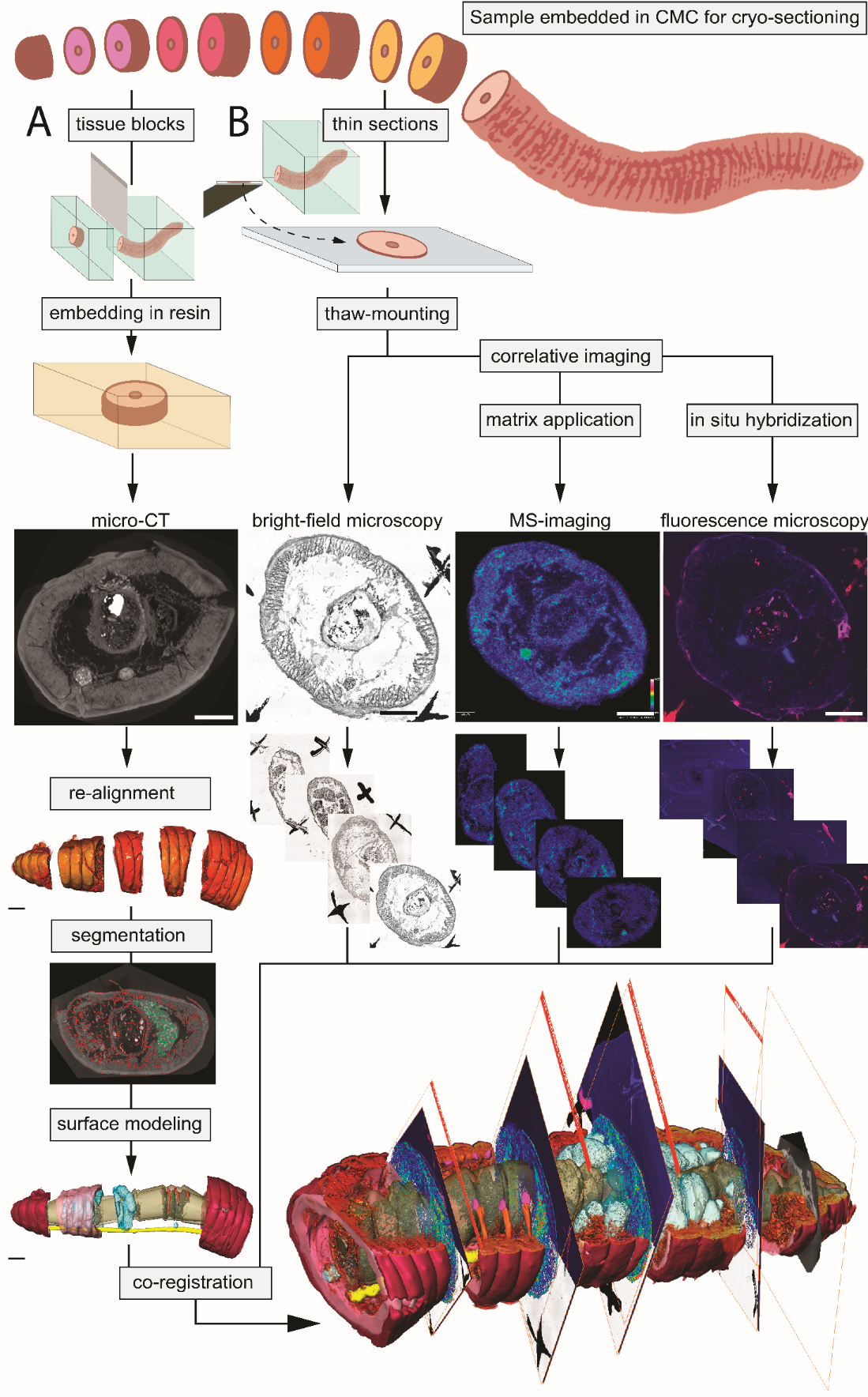

**Figure S-1: Schematic workflow of 3D anatomy-guided multimodal imaging of an earthworm** (from end to front, to display the correct order of sample and image acquisition)**.** The frozen embedded sample was physically divided into tissue blocks with a razorblade **(A)**, from which thin-sections were taken with a cryotome **(B)**. The tissue blocks (*n* = 5) were contrasted and scanned with microCT. The microCT datasets were realigned in 3D space and served as a template for threshold-based organ segmentation and surface modeling. Thin-sections were thaw-mounted onto glass slides and imaged with bright-field microscopy, MALDI-MSI for spatial metabolite distributions followed by fluorescence microscopy. Bright-field microscopy, MALDI-MSI and fluorescence microscopy datasets were co-registered into the spaces between the individual microCT datasets. Scale bars of 2D imaging modalities, 500 µm and of 3D models, 2 mm.

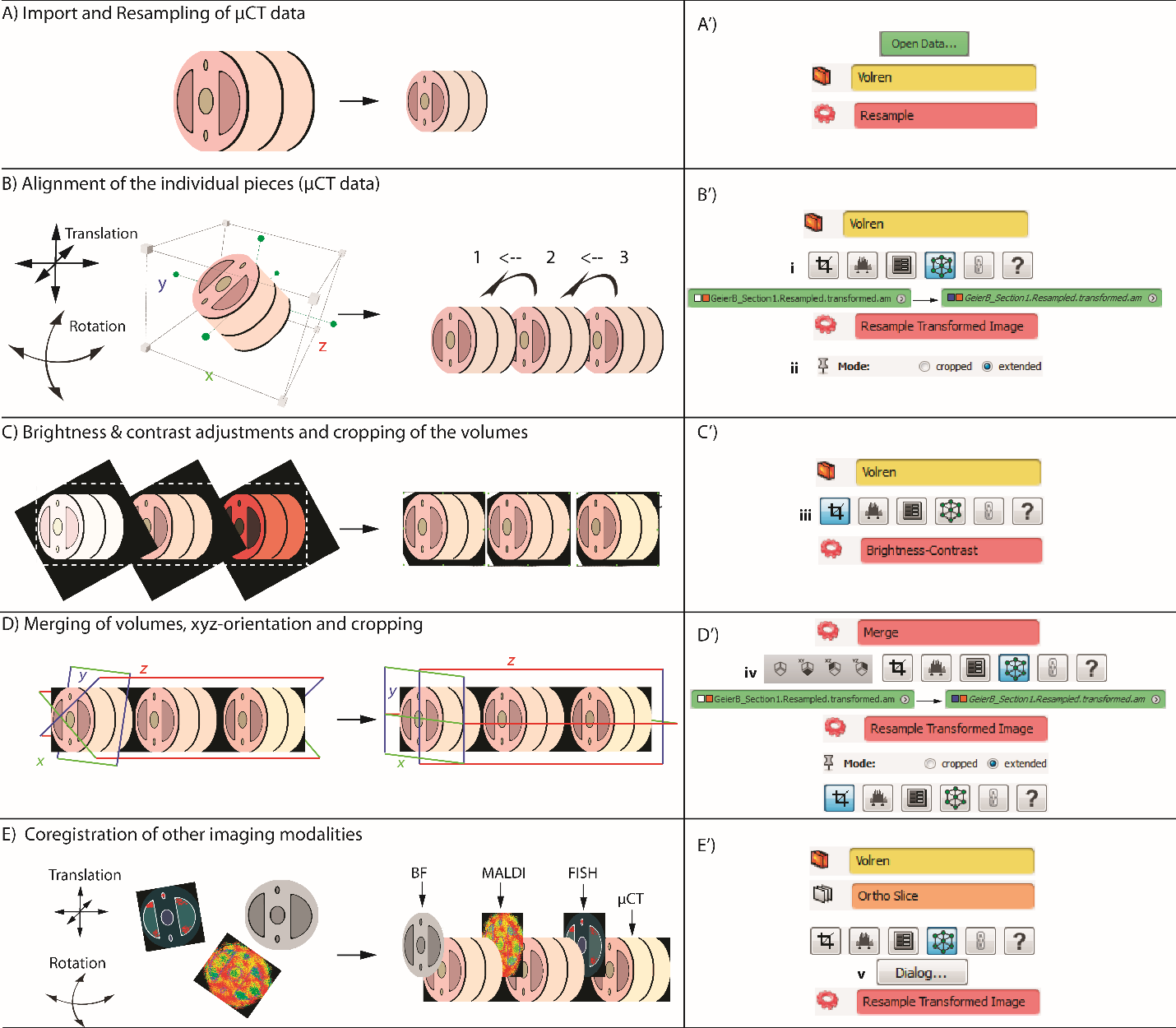

Figure S-2: Scheme of the general registration workflow (left, A–E) and screenshots of the basic modules in AMIRA (right, A’–E’). Workflow: A) Import and resampling; B) 3D Alignment (co-registration without overlapping) of μCT volumes; C) Adjustment of brightness and contrast and cropping (white dashed line) of individual volumes; D) Merging of volumes including blending (faster with cropped volumes), *xyz*-global orientation and second cropping of the final volume; E) Integration of the imaging data of the other modalities (BF, FISH, MALDI). Screen shots: red, computing modules (Resample, Resample Transformed Image, Brightness-Contrast, Merge); orange, 2D display module (Ortho Slice); yellow, 3D display module (Volren); green, datasets and general commands (open, apply etc.); i, editors with transform editor highlighted; ii, mode (extended) to apply a transformation; iii, editors with crop editor highlighted; iv, viewer orientation module to reset the perspective (left to right: center dataset, *xy*, *xz*, *yz*); v, dialog with transformation parameters.

|  |  |  |
| --- | --- | --- |
| 99.7925 | 0.230815 | 0.230815 |
| 99.9407 | 0.230815 | 0.230815 |
| 100.089 | 0.230815 | 0.230815 |
| 100.237 | 0.106716 | 0.106716 |
| 100.386 | 0.0914585 | 0.0914585 |
| 100.534 | 0.0874226 | 0.0874226 |
| 100.682 | 0.0843534 | 0.0843534 |
| 100.83 | 0.0820178 | 0.0820178 |
| 100.979 | 0.0996448 | 0.0996448 |
| 101.127 | 0.104282 | 0.104282 |
| 101.275 | 0.0795611 | 0.0795611 |
| 101.423 | 0.0765022 | 0.0765022 |
| 101.572 | 0.0745678 | 0.0745678 |
| 101.72 | 0.0725595 | 0.0725595 |
| 101.868 | 0.0715597 | 0.0715597 |
| 102.016 | 0.293388 | 0.293388 |
| 102.165 | 0.131894 | 0.131894 |
| 102.313 | 0.0729261 | 0.0729261 |
| 102.461 | 0.0678944 | 0.0678944 |
| 102.609 | 0.0655446 | 0.0655446 |
| 102.758 | 0.0643608 | 0.0643608 |
| 102.906 | 0.0662781 | 0.0662781 |
| 103.054 | 0.0781915 | 0.0781915 |
| 103.203 | 0.062477 | 0.062477 |
| 103.351 | 0.0595076 | 0.0595076 |
| 103.499 | 0.0587683 | 0.0587683 |
| 103.647 | 0.0569658 | 0.0569658 |
| 103.796 | 0.0716368 | 0.0716368 |
| 103.944 | 3.25681 | 3.25681 |
| 104.092 | 23.1879 | 23.1879 |
| 104.24 | 4.46506 | 4.46506 |
| 104.389 | 1.032 | 1.032 |
| 104.537 | 0.659712 | 0.659712 |
| 104.685 | 0.479728 | 0.479728 |
| 104.833 | 0.421989 | 0.421989 |
| 104.982 | 0.843442 | 0.843442 |
| 105.13 | 1.73281 | 1.73281 |
| 105.278 | 0.458623 | 0.458623 |
| 105.427 | 0.329525 | 0.329525 |
| 105.575 | 0.403162 | 0.403162 |
| 105.723 | 0.266919 | 0.266919 |
| 105.871 | 0.253966 | 0.253966 |
| 106.02 | 0.298917 | 0.298917 |
| 106.168 | 0.275459 | 0.275459 |
| 106.316 | 0.221663 | 0.221663 |
| 106.464 | 0.209936 | 0.209936 |
| 106.613 | 0.203585 | 0.203585 |
| 106.761 | 0.194756 | 0.194756 |
| 106.909 | 0.190158 | 0.190158 |
| 107.057 | 0.194394 | 0.194394 |
| 107.206 | 0.178717 | 0.178717 |
| 107.354 | 0.172173 | 0.172173 |
| 107.502 | 0.166433 | 0.166433 |
| 107.65 | 0.160946 | 0.160946 |
| 107.799 | 0.15639 | 0.15639 |
| 107.947 | 0.157065 | 0.157065 |
| 108.095 | 0.15734 | 0.15734 |
| 108.244 | 0.145308 | 0.145308 |
| 108.392 | 0.140895 | 0.140895 |
| 108.54 | 0.13668 | 0.13668 |
| 108.688 | 0.133575 | 0.133575 |
| 108.837 | 0.130784 | 0.130784 |
| 108.985 | 0.133469 | 0.133469 |
| 109.133 | 0.13255 | 0.13255 |
| 109.281 | 0.123245 | 0.123245 |
| 109.43 | 0.119129 | 0.119129 |
| 109.578 | 0.116033 | 0.116033 |
| 109.726 | 0.113511 | 0.113511 |
| 109.874 | 0.110949 | 0.110949 |
| 110.023 | 0.120805 | 0.120805 |
| 110.171 | 0.111659 | 0.111659 |
| 110.319 | 0.105309 | 0.105309 |
| 110.468 | 0.103927 | 0.103927 |
| 110.616 | 0.101705 | 0.101705 |
| 110.764 | 0.0982015 | 0.0982015 |
| 110.912 | 0.0980681 | 0.0980681 |
| 111.061 | 0.109011 | 0.109011 |
| 111.209 | 0.0964008 | 0.0964008 |
| 111.357 | 0.0917461 | 0.0917461 |
| 111.505 | 0.0892314 | 0.0892314 |
| 111.654 | 0.087537 | 0.087537 |
| 111.802 | 0.0861118 | 0.0861118 |
| 111.95 | 0.118428 | 0.118428 |
| 112.098 | 0.11541 | 0.11541 |
| 112.247 | 0.0895791 | 0.0895791 |
| 112.395 | 0.0855592 | 0.0855592 |
| 112.543 | 0.0864941 | 0.0864941 |
| 112.691 | 0.091587 | 0.091587 |
| 112.84 | 0.115564 | 0.115564 |
| 112.988 | 0.133704 | 0.133704 |
| 113.136 | 0.138548 | 0.138548 |
| 113.285 | 0.12885 | 0.12885 |
| 113.433 | 0.114711 | 0.114711 |
| 113.581 | 0.0987111 | 0.0987111 |
| 113.729 | 0.0865922 | 0.0865922 |
| 113.878 | 0.0804289 | 0.0804289 |
| 114.026 | 0.0866759 | 0.0866759 |
| 114.174 | 0.086003 | 0.086003 |
| 114.322 | 0.0724173 | 0.0724173 |
| 114.471 | 0.0690109 | 0.0690109 |
| 114.619 | 0.067329 | 0.067329 |
| 114.767 | 0.0661215 | 0.0661215 |
| 114.915 | 0.0684023 | 0.0684023 |
| 115.064 | 0.0684705 | 0.0684705 |
| 115.212 | 0.0636516 | 0.0636516 |
| 115.36 | 0.0611067 | 0.0611067 |
| 115.509 | 0.059321 | 0.059321 |
| 115.657 | 0.0576686 | 0.0576686 |
| 115.805 | 0.0561111 | 0.0561111 |
| 115.953 | 0.0601072 | 0.0601072 |
| 116.102 | 0.0818998 | 0.0818998 |
| 116.25 | 0.057367 | 0.057367 |
| 116.398 | 0.0530725 | 0.0530725 |
| 116.546 | 0.051909 | 0.051909 |
| 116.695 | 0.0508046 | 0.0508046 |
| 116.843 | 0.0512511 | 0.0512511 |
| 116.991 | 0.0548001 | 0.0548001 |
| 117.139 | 0.0555114 | 0.0555114 |
| 117.288 | 0.0489186 | 0.0489186 |
| 117.436 | 0.0473965 | 0.0473965 |
| 117.584 | 0.0464653 | 0.0464653 |
| 117.732 | 0.0454889 | 0.0454889 |
| 117.881 | 0.0452728 | 0.0452728 |
| 118.029 | 0.367404 | 0.367404 |
| 118.177 | 0.210134 | 0.210134 |
| 118.326 | 0.0557299 | 0.0557299 |
| 118.474 | 0.0457149 | 0.0457149 |
| 118.622 | 0.0439129 | 0.0439129 |
| 118.77 | 0.0427811 | 0.0427811 |
| 118.919 | 0.0435021 | 0.0435021 |
| 119.067 | 0.0671547 | 0.0671547 |
| 119.215 | 0.0462249 | 0.0462249 |
| 119.363 | 0.0404993 | 0.0404993 |
| 119.512 | 0.0399421 | 0.0399421 |
| 119.66 | 0.0392977 | 0.0392977 |
| 119.808 | 0.0393459 | 0.0393459 |
| 119.956 | 0.0474376 | 0.0474376 |
| 120.105 | 0.0669377 | 0.0669377 |
| 120.253 | 0.0413847 | 0.0413847 |
| 120.401 | 0.0367853 | 0.0367853 |
| 120.549 | 0.0357403 | 0.0357403 |
| 120.698 | 0.0351409 | 0.0351409 |
| 120.846 | 0.0407669 | 0.0407669 |
| 120.994 | 0.0651204 | 0.0651204 |
| 121.143 | 0.0435511 | 0.0435511 |
| 121.291 | 0.0360824 | 0.0360824 |
| 121.439 | 0.0346668 | 0.0346668 |
| 121.587 | 0.0344429 | 0.0344429 |
| 121.736 | 0.0416015 | 0.0416015 |
| 121.884 | 0.0384136 | 0.0384136 |
| 122.032 | 0.048182 | 0.048182 |
| 122.18 | 0.0375645 | 0.0375645 |
| 122.329 | 0.0318137 | 0.0318137 |
| 122.477 | 0.0314232 | 0.0314232 |
| 122.625 | 0.0307242 | 0.0307242 |
| 122.773 | 0.030325 | 0.030325 |
| 122.922 | 0.0324889 | 0.0324889 |
| 123.07 | 0.0392852 | 0.0392852 |
| 123.218 | 0.0319253 | 0.0319253 |
| 123.367 | 0.0295704 | 0.0295704 |
| 123.515 | 0.0290511 | 0.0290511 |
| 123.663 | 0.0286942 | 0.0286942 |
| 123.811 | 0.028594 | 0.028594 |
| 123.96 | 0.0319181 | 0.0319181 |
| 124.108 | 0.0384268 | 0.0384268 |
| 124.256 | 0.0287404 | 0.0287404 |
| 124.404 | 0.0273384 | 0.0273384 |
| 124.553 | 0.0277605 | 0.0277605 |
| 124.701 | 0.026914 | 0.026914 |
| 124.849 | 0.0384525 | 0.0384525 |
| 124.997 | 0.120604 | 0.120604 |
| 125.146 | 0.0584889 | 0.0584889 |
| 125.294 | 0.0376129 | 0.0376129 |
| 125.442 | 0.0336912 | 0.0336912 |
| 125.59 | 0.0318941 | 0.0318941 |
| 125.739 | 0.0308701 | 0.0308701 |
| 125.887 | 0.0315905 | 0.0315905 |
| 126.035 | 0.041383 | 0.041383 |
| 126.184 | 0.040021 | 0.040021 |
| 126.332 | 0.0282394 | 0.0282394 |
| 126.48 | 0.0267547 | 0.0267547 |
| 126.628 | 0.0260355 | 0.0260355 |
| 126.777 | 0.0255454 | 0.0255454 |
| 126.925 | 0.0264276 | 0.0264276 |
| 127.073 | 0.0456895 | 0.0456895 |
| 127.221 | 0.0373543 | 0.0373543 |
| 127.37 | 0.0255785 | 0.0255785 |
| 127.518 | 0.0245936 | 0.0245936 |
| 127.666 | 0.0239804 | 0.0239804 |
| 127.814 | 0.0237642 | 0.0237642 |
| 127.963 | 0.0615312 | 0.0615312 |
| 128.111 | 0.0553256 | 0.0553256 |
| 128.259 | 0.0337478 | 0.0337478 |
| 128.408 | 0.0262072 | 0.0262072 |
| 128.556 | 0.026165 | 0.026165 |
| 128.704 | 0.0244776 | 0.0244776 |
| 128.852 | 0.0235662 | 0.0235662 |
| 129.001 | 0.0286246 | 0.0286246 |
| 129.149 | 0.0566832 | 0.0566832 |
| 129.297 | 0.0306626 | 0.0306626 |

|  |  |
| --- | --- |
| 99.7925 | 4.69946 |
| 99.9407 | 4.69946 |
| 100.089 | 4.69946 |
| 100.237 | 2.98785 |
| 100.386 | 2.67587 |
| 100.534 | 2.56446 |
| 100.682 | 2.49606 |
| 100.83 | 2.42941 |
| 100.979 | 2.59017 |
| 101.127 | 2.66045 |
| 101.275 | 2.31635 |
| 101.423 | 2.25164 |
| 101.572 | 2.18633 |
| 101.72 | 2.11967 |
| 101.868 | 2.12145 |
| 102.016 | 4.91541 |
| 102.165 | 3.31604 |
| 102.313 | 2.08309 |
| 102.461 | 1.95547 |
| 102.609 | 1.89491 |
| 102.758 | 1.84091 |
| 102.906 | 1.85799 |
| 103.054 | 2.0065 |
| 103.203 | 1.77486 |
| 103.351 | 1.70775 |
| 103.499 | 1.68761 |
| 103.647 | 1.63484 |
| 103.796 | 1.90097 |
| 103.944 | 65.7269 |
| 104.092 | 446.186 |
| 104.24 | 135.515 |
| 104.389 | 31.8726 |
| 104.537 | 19.1207 |
| 104.685 | 13.769 |
| 104.833 | 12.219 |
| 104.982 | 18.3144 |
| 105.13 | 43.0068 |
| 105.278 | 13.8412 |
| 105.427 | 8.84221 |
| 105.575 | 11.2455 |
| 105.723 | 7.60815 |
| 105.871 | 7.0482 |
| 106.02 | 7.71262 |
| 106.168 | 7.67448 |
| 106.316 | 6.19848 |
| 106.464 | 5.80169 |
| 106.613 | 5.62152 |
| 106.761 | 5.34856 |
| 106.909 | 5.21887 |
| 107.057 | 5.389 |
| 107.206 | 4.91486 |
| 107.354 | 4.68769 |
| 107.502 | 4.52264 |
| 107.65 | 4.35853 |
| 107.799 | 4.22807 |
| 107.947 | 4.15747 |
| 108.095 | 4.13703 |
| 108.244 | 3.87932 |
| 108.392 | 3.75706 |
| 108.54 | 3.64754 |
| 108.688 | 3.54489 |
| 108.837 | 3.4583 |
| 108.985 | 3.50533 |
| 109.133 | 3.65779 |
| 109.281 | 3.28441 |
| 109.43 | 3.16482 |
| 109.578 | 3.07549 |
| 109.726 | 2.99569 |
| 109.874 | 2.92445 |
| 110.023 | 3.18603 |
| 110.171 | 2.93877 |
| 110.319 | 2.75262 |
| 110.468 | 2.68023 |
| 110.616 | 2.61251 |
| 110.764 | 2.54095 |
| 110.912 | 2.5148 |
| 111.061 | 2.77633 |
| 111.209 | 2.53568 |
| 111.357 | 2.38912 |
| 111.505 | 2.32619 |
| 111.654 | 2.27824 |
| 111.802 | 2.22144 |
| 111.95 | 2.57324 |
| 112.098 | 2.68329 |
| 112.247 | 2.25272 |
| 112.395 | 2.19999 |
| 112.543 | 2.27259 |
| 112.691 | 2.49313 |
| 112.84 | 3.22225 |
| 112.988 | 3.97194 |
| 113.136 | 4.3604 |
| 113.285 | 4.10961 |
| 113.433 | 3.57421 |
| 113.581 | 2.95491 |
| 113.729 | 2.47999 |
| 113.878 | 2.21979 |
| 114.026 | 2.1823 |
| 114.174 | 2.11474 |
| 114.322 | 1.8873 |
| 114.471 | 1.79933 |
| 114.619 | 1.74096 |
| 114.767 | 1.69325 |
| 114.915 | 1.70831 |
| 115.064 | 1.73386 |
| 115.212 | 1.62196 |
| 115.36 | 1.56297 |
| 115.509 | 1.52609 |
| 115.657 | 1.4838 |
| 115.805 | 1.44124 |
| 115.953 | 1.48877 |
| 116.102 | 1.72667 |
| 116.25 | 1.4359 |
| 116.398 | 1.34955 |
| 116.546 | 1.31777 |
| 116.695 | 1.28647 |
| 116.843 | 1.27853 |
| 116.991 | 1.32362 |
| 117.139 | 1.30743 |
| 117.288 | 1.21587 |
| 117.436 | 1.18263 |
| 117.584 | 1.15934 |
| 117.732 | 1.13657 |
| 117.881 | 1.13493 |
| 118.029 | 3.1166 |
| 118.177 | 3.01533 |
| 118.326 | 1.27909 |
| 118.474 | 1.11909 |
| 118.622 | 1.07646 |
| 118.77 | 1.05288 |
| 118.919 | 1.08211 |
| 119.067 | 1.30116 |
| 119.215 | 1.10457 |
| 119.363 | 1.0016 |
| 119.512 | 0.975718 |
| 119.66 | 0.958301 |
| 119.808 | 0.97272 |
| 119.956 | 1.13198 |
| 120.105 | 1.63292 |
| 120.253 | 1.07013 |
| 120.401 | 0.916758 |
| 120.549 | 0.887318 |
| 120.698 | 0.864394 |
| 120.846 | 0.943914 |
| 120.994 | 1.44718 |
| 121.143 | 1.13974 |
| 121.291 | 0.907977 |
| 121.439 | 0.861796 |
| 121.587 | 0.846308 |
| 121.736 | 0.899342 |
| 121.884 | 0.886615 |
| 122.032 | 0.991913 |
| 122.18 | 0.884926 |
| 122.329 | 0.779486 |
| 122.477 | 0.779414 |
| 122.625 | 0.767567 |
| 122.773 | 0.747272 |
| 122.922 | 0.783463 |
| 123.07 | 0.992599 |
| 123.218 | 0.827975 |
| 123.367 | 0.747856 |
| 123.515 | 0.73422 |
| 123.663 | 0.716959 |
| 123.811 | 0.702412 |
| 123.96 | 0.747482 |
| 124.108 | 0.871587 |
| 124.256 | 0.710672 |
| 124.404 | 0.685404 |
| 124.553 | 0.722744 |
| 124.701 | 0.684942 |
| 124.849 | 0.943369 |
| 124.997 | 4.55569 |
| 125.146 | 2.16203 |
| 125.294 | 1.15176 |
| 125.442 | 0.98997 |
| 125.59 | 0.919232 |
| 125.739 | 0.864366 |
| 125.887 | 0.834616 |
| 126.035 | 0.986116 |
| 126.184 | 0.896623 |
| 126.332 | 0.719396 |
| 126.48 | 0.685283 |
| 126.628 | 0.665657 |
| 126.777 | 0.643857 |
| 126.925 | 0.661065 |
| 127.073 | 0.880474 |
| 127.221 | 0.795016 |
| 127.37 | 0.639787 |
| 127.518 | 0.614581 |
| 127.666 | 0.603351 |
| 127.814 | 0.593817 |
| 127.963 | 1.07905 |
| 128.111 | 0.898638 |
| 128.259 | 0.682753 |
| 128.408 | 0.595153 |
| 128.556 | 0.589333 |
| 128.704 | 0.567241 |
| 128.852 | 0.551466 |
| 129.001 | 0.644197 |
| 129.149 | 1.00696 |
| 129.297 | 0.657971 |

|  |  |
| --- | --- |
| 99.7851 | 0.769973 |
| 99.9333 | 0.769973 |
| 100.082 | 0.769973 |
| 100.23 | 0.300264 |
| 100.378 | 0.221989 |
| 100.526 | 0.206416 |
| 100.675 | 0.197053 |
| 100.823 | 0.191433 |
| 100.971 | 0.236044 |
| 101.119 | 0.25531 |
| 101.268 | 0.186489 |
| 101.416 | 0.176622 |
| 101.564 | 0.170858 |
| 101.712 | 0.164591 |
| 101.861 | 0.163531 |
| 102.009 | 0.396616 |
| 102.157 | 0.247126 |
| 102.305 | 0.15975 |
| 102.454 | 0.149953 |
| 102.602 | 0.144825 |
| 102.75 | 0.140955 |
| 102.898 | 0.144681 |
| 103.047 | 0.158686 |
| 103.195 | 0.137463 |
| 103.343 | 0.130778 |
| 103.491 | 0.129048 |
| 103.64 | 0.124727 |
| 103.788 | 0.166336 |
| 103.936 | 6.26603 |
| 104.084 | 30.6901 |
| 104.233 | 8.91709 |
| 104.381 | 2.10688 |
| 104.529 | 1.30975 |
| 104.677 | 0.966397 |
| 104.826 | 0.858641 |
| 104.974 | 1.52259 |
| 105.122 | 2.65283 |
| 105.27 | 0.956678 |
| 105.419 | 0.624396 |
| 105.567 | 0.737161 |
| 105.715 | 0.543219 |
| 105.863 | 0.515049 |
| 106.012 | 0.613178 |
| 106.16 | 0.556073 |
| 106.308 | 0.449653 |
| 106.456 | 0.420582 |
| 106.605 | 0.406568 |
| 106.753 | 0.387201 |
| 106.901 | 0.384154 |
| 107.049 | 0.410455 |
| 107.198 | 0.360905 |
| 107.346 | 0.341003 |
| 107.494 | 0.329042 |
| 107.642 | 0.316266 |
| 107.791 | 0.306391 |
| 107.939 | 0.323904 |
| 108.087 | 0.334063 |
| 108.235 | 0.286805 |
| 108.384 | 0.273257 |
| 108.532 | 0.264504 |
| 108.68 | 0.256307 |
| 108.828 | 0.251002 |
| 108.977 | 0.268494 |
| 109.125 | 0.274611 |
| 109.273 | 0.238021 |
| 109.421 | 0.227765 |
| 109.57 | 0.220869 |
| 109.718 | 0.214279 |
| 109.866 | 0.209438 |
| 110.014 | 0.253067 |
| 110.163 | 0.222076 |
| 110.311 | 0.198812 |
| 110.459 | 0.194287 |
| 110.607 | 0.188963 |
| 110.756 | 0.181942 |
| 110.904 | 0.182162 |
| 111.052 | 0.220195 |
| 111.2 | 0.186093 |
| 111.349 | 0.169298 |
| 111.497 | 0.164252 |
| 111.645 | 0.159295 |
| 111.793 | 0.157099 |
| 111.942 | 0.276784 |
| 112.09 | 0.27107 |
| 112.238 | 0.182116 |
| 112.386 | 0.165196 |
| 112.535 | 0.165123 |
| 112.683 | 0.178432 |
| 112.831 | 0.240448 |
| 112.979 | 0.292811 |
| 113.128 | 0.320172 |
| 113.276 | 0.303377 |
| 113.424 | 0.267125 |
| 113.572 | 0.218428 |
| 113.721 | 0.178897 |
| 113.869 | 0.158098 |
| 114.017 | 0.165888 |
| 114.165 | 0.16113 |
| 114.314 | 0.135316 |
| 114.462 | 0.127312 |
| 114.61 | 0.122367 |
| 114.758 | 0.120395 |
| 114.907 | 0.125356 |
| 115.055 | 0.128 |
| 115.203 | 0.115973 |
| 115.351 | 0.109963 |
| 115.5 | 0.105462 |
| 115.648 | 0.100701 |
| 115.796 | 0.09675 |
| 115.944 | 0.11585 |
| 116.093 | 0.167294 |
| 116.241 | 0.107041 |
| 116.389 | 0.0917523 |
| 116.537 | 0.0879301 |
| 116.686 | 0.0853496 |
| 116.834 | 0.0860632 |
| 116.982 | 0.0979709 |
| 117.13 | 0.0965471 |
| 117.279 | 0.0824963 |
| 117.427 | 0.0788616 |
| 117.575 | 0.076867 |
| 117.723 | 0.0749284 |
| 117.872 | 0.0793434 |
| 118.02 | 0.818677 |
| 118.168 | 0.49649 |
| 118.316 | 0.111367 |
| 118.465 | 0.0780427 |
| 118.613 | 0.0731828 |
| 118.761 | 0.0705351 |
| 118.909 | 0.0792565 |
| 119.058 | 0.134215 |
| 119.206 | 0.0862062 |
| 119.354 | 0.0689233 |
| 119.502 | 0.0664111 |
| 119.651 | 0.0652822 |
| 119.799 | 0.065969 |
| 119.947 | 0.0828772 |
| 120.095 | 0.104938 |
| 120.244 | 0.0701491 |
| 120.392 | 0.0616353 |
| 120.54 | 0.059598 |
| 120.688 | 0.0582218 |
| 120.837 | 0.077284 |
| 120.985 | 0.165098 |
| 121.133 | 0.0954546 |
| 121.281 | 0.0679337 |
| 121.43 | 0.0626722 |
| 121.578 | 0.0626805 |
| 121.726 | 0.0765281 |
| 121.874 | 0.0673702 |
| 122.023 | 0.0967859 |
| 122.171 | 0.0769947 |
| 122.319 | 0.0570756 |
| 122.467 | 0.056358 |
| 122.616 | 0.0546293 |
| 122.764 | 0.0528231 |
| 122.912 | 0.057883 |
| 123.06 | 0.0885243 |
| 123.209 | 0.0647323 |
| 123.357 | 0.0538744 |
| 123.505 | 0.052069 |
| 123.653 | 0.0508284 |
| 123.802 | 0.0500932 |
| 123.95 | 0.0647176 |
| 124.098 | 0.0837539 |
| 124.246 | 0.0550927 |
| 124.395 | 0.0512405 |
| 124.543 | 0.0549181 |
| 124.691 | 0.0501176 |
| 124.839 | 0.157111 |
| 124.988 | 0.510877 |
| 125.136 | 0.205753 |
| 125.284 | 0.0995998 |
| 125.432 | 0.0833508 |
| 125.581 | 0.076059 |
| 125.729 | 0.0703294 |
| 125.877 | 0.0706196 |
| 126.025 | 0.103392 |
| 126.174 | 0.0875306 |
| 126.322 | 0.0589212 |
| 126.47 | 0.0542984 |
| 126.618 | 0.0520365 |
| 126.767 | 0.050243 |
| 126.915 | 0.0532271 |
| 127.063 | 0.0925128 |
| 127.211 | 0.0783981 |
| 127.36 | 0.0513402 |
| 127.508 | 0.0478918 |
| 127.656 | 0.0463109 |
| 127.804 | 0.0487753 |
| 127.953 | 0.153973 |
| 128.101 | 0.105512 |
| 128.249 | 0.0656177 |
| 128.397 | 0.0521034 |
| 128.546 | 0.053002 |
| 128.694 | 0.0486683 |
| 128.842 | 0.0462567 |
| 128.99 | 0.0645962 |
| 129.139 | 0.164987 |
| 129.287 | 0.0855009 |

|  |  |
| --- | --- |
| 99.7798 | 4.00208 |
| 99.9281 | 4.00208 |
| 100.076 | 4.00208 |
| 100.225 | 1.39642 |
| 100.373 | 1.27972 |
| 100.521 | 1.23041 |
| 100.669 | 1.1881 |
| 100.818 | 1.16007 |
| 100.966 | 1.32368 |
| 101.114 | 1.21828 |
| 101.262 | 1.08907 |
| 101.411 | 1.0683 |
| 101.559 | 1.04316 |
| 101.707 | 1.00499 |
| 101.855 | 0.988572 |
| 102.004 | 1.54019 |
| 102.152 | 1.01909 |
| 102.3 | 0.928763 |
| 102.449 | 0.903704 |
| 102.597 | 0.885219 |
| 102.745 | 0.864186 |
| 102.893 | 0.879342 |
| 103.042 | 0.900882 |
| 103.19 | 0.817271 |
| 103.338 | 0.804477 |
| 103.486 | 0.792028 |
| 103.635 | 0.772477 |
| 103.783 | 1.02364 |
| 103.931 | 34.9902 |
| 104.079 | 231.873 |
| 104.228 | 13.736 |
| 104.376 | 4.00178 |
| 104.524 | 3.00934 |
| 104.672 | 2.50113 |
| 104.821 | 2.30292 |
| 104.969 | 6.68288 |
| 105.117 | 7.52917 |
| 105.266 | 2.18185 |
| 105.414 | 1.96569 |
| 105.562 | 1.91002 |
| 105.71 | 1.68571 |
| 105.859 | 1.68304 |
| 106.007 | 2.37976 |
| 106.155 | 1.69177 |
| 106.303 | 1.47552 |
| 106.452 | 1.41037 |
| 106.6 | 1.36732 |
| 106.748 | 1.3156 |
| 106.896 | 1.35375 |
| 107.045 | 1.48955 |
| 107.193 | 1.25347 |
| 107.341 | 1.20708 |
| 107.489 | 1.16693 |
| 107.638 | 1.131 |
| 107.786 | 1.10424 |
| 107.934 | 1.21314 |
| 108.083 | 1.16875 |
| 108.231 | 1.03494 |
| 108.379 | 1.00268 |
| 108.527 | 0.978679 |
| 108.676 | 0.950544 |
| 108.824 | 0.947341 |
| 108.972 | 1.11275 |
| 109.12 | 1.0662 |
| 109.269 | 0.909696 |
| 109.417 | 0.87947 |
| 109.565 | 0.851731 |
| 109.713 | 0.828092 |
| 109.862 | 0.816047 |
| 110.01 | 0.986139 |
| 110.158 | 0.812994 |
| 110.306 | 0.770696 |
| 110.455 | 0.76793 |
| 110.603 | 0.741678 |
| 110.751 | 0.717849 |
| 110.899 | 0.727379 |
| 111.048 | 0.963262 |
| 111.196 | 0.733663 |
| 111.344 | 0.688398 |
| 111.493 | 0.674337 |
| 111.641 | 0.656236 |
| 111.789 | 0.645312 |
| 111.937 | 1.20001 |
| 112.086 | 0.95378 |
| 112.234 | 0.691212 |
| 112.382 | 0.685049 |
| 112.53 | 0.722467 |
| 112.679 | 0.847441 |
| 112.827 | 1.35285 |
| 112.975 | 1.54136 |
| 113.123 | 1.55864 |
| 113.272 | 1.37643 |
| 113.42 | 1.11813 |
| 113.568 | 0.883129 |
| 113.716 | 0.734106 |
| 113.865 | 0.669521 |
| 114.013 | 0.747263 |
| 114.161 | 0.652064 |
| 114.31 | 0.568679 |
| 114.458 | 0.541215 |
| 114.606 | 0.521085 |
| 114.754 | 0.525068 |
| 114.903 | 0.566005 |
| 115.051 | 0.549637 |
| 115.199 | 0.483493 |
| 115.347 | 0.470888 |
| 115.496 | 0.455051 |
| 115.644 | 0.441036 |
| 115.792 | 0.428449 |
| 115.94 | 0.50828 |
| 116.089 | 0.704206 |
| 116.237 | 0.430898 |
| 116.385 | 0.410772 |
| 116.533 | 0.396826 |
| 116.682 | 0.392198 |
| 116.83 | 0.40282 |
| 116.978 | 0.454806 |
| 117.127 | 0.424967 |
| 117.275 | 0.377446 |
| 117.423 | 0.369117 |
| 117.571 | 0.361901 |
| 117.72 | 0.353458 |
| 117.868 | 0.35031 |
| 118.016 | 5.34907 |
| 118.164 | 1.54881 |
| 118.313 | 0.391581 |
| 118.461 | 0.352071 |
| 118.609 | 0.34028 |
| 118.757 | 0.334991 |
| 118.906 | 0.360575 |
| 119.054 | 0.731408 |
| 119.202 | 0.364087 |
| 119.35 | 0.330551 |
| 119.499 | 0.322643 |
| 119.647 | 0.31806 |
| 119.795 | 0.333435 |
| 119.944 | 0.410439 |
| 120.092 | 0.437916 |
| 120.24 | 0.309744 |
| 120.388 | 0.30045 |
| 120.537 | 0.296143 |
| 120.685 | 0.291129 |
| 120.833 | 0.416851 |
| 120.981 | 0.74306 |
| 121.13 | 0.418812 |
| 121.278 | 0.326891 |
| 121.426 | 0.313602 |
| 121.574 | 0.313447 |
| 121.723 | 0.419105 |
| 121.871 | 0.350999 |
| 122.019 | 0.51293 |
| 122.167 | 0.325454 |
| 122.316 | 0.287417 |
| 122.464 | 0.291958 |
| 122.612 | 0.284709 |
| 122.761 | 0.281857 |
| 122.909 | 0.314795 |
| 123.057 | 0.469637 |
| 123.205 | 0.303264 |
| 123.354 | 0.283474 |
| 123.502 | 0.278877 |
| 123.65 | 0.273526 |
| 123.798 | 0.271288 |
| 123.947 | 0.362226 |
| 124.095 | 0.365371 |
| 124.243 | 0.265312 |
| 124.391 | 0.268727 |
| 124.54 | 0.290365 |
| 124.688 | 0.26505 |
| 124.836 | 0.824231 |
| 124.984 | 2.37341 |
| 125.133 | 0.676618 |
| 125.281 | 0.443573 |
| 125.429 | 0.396335 |
| 125.577 | 0.379965 |
| 125.726 | 0.359923 |
| 125.874 | 0.368525 |
| 126.022 | 0.502209 |
| 126.171 | 0.347735 |
| 126.319 | 0.294774 |
| 126.467 | 0.285377 |
| 126.615 | 0.275134 |
| 126.764 | 0.267119 |
| 126.912 | 0.289892 |
| 127.06 | 0.546651 |
| 127.208 | 0.329589 |
| 127.357 | 0.268017 |
| 127.505 | 0.260582 |
| 127.653 | 0.254509 |
| 127.801 | 0.261643 |
| 127.95 | 0.781839 |
| 128.098 | 0.468259 |
| 128.246 | 0.296868 |
| 128.394 | 0.267693 |
| 128.543 | 0.281215 |
| 128.691 | 0.255794 |
| 128.839 | 0.251376 |
| 128.988 | 0.329691 |
| 129.136 | 0.603403 |
| 129.284 | 0.290771 |

**Figure S-3: Total ion spectra of each tissue section (sections s1–s4) analyzed with MALDI-TOF-MSI (*m/z* 100–1200 Da, *y*-axis displays relative abundance).**

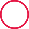

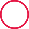

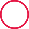

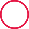

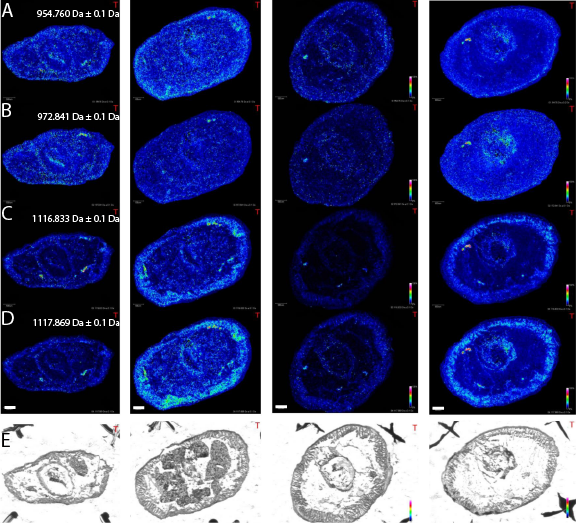
**­**

Figure S-4: Distributions of three different metabolites (A–D) found by ROI matching with bacterial regions, detected with FISH (encircled in the bright-field image of E). Sections s4 to s1 from left to right. Ion maps of *m/z* 954.760 (A), *m/z* 972.841 (B), *m/z* 1116.833 (C), and *m/z* 1117.869 (^13^C isotope of metabolite *m/z* 1116.833) (D), are displayed as heat maps ± 100 mDa. (E) Brightfield images are labeled with the bacteria-containing regions, detected with FISH and encircled in red. Scale bar: 500 µm.

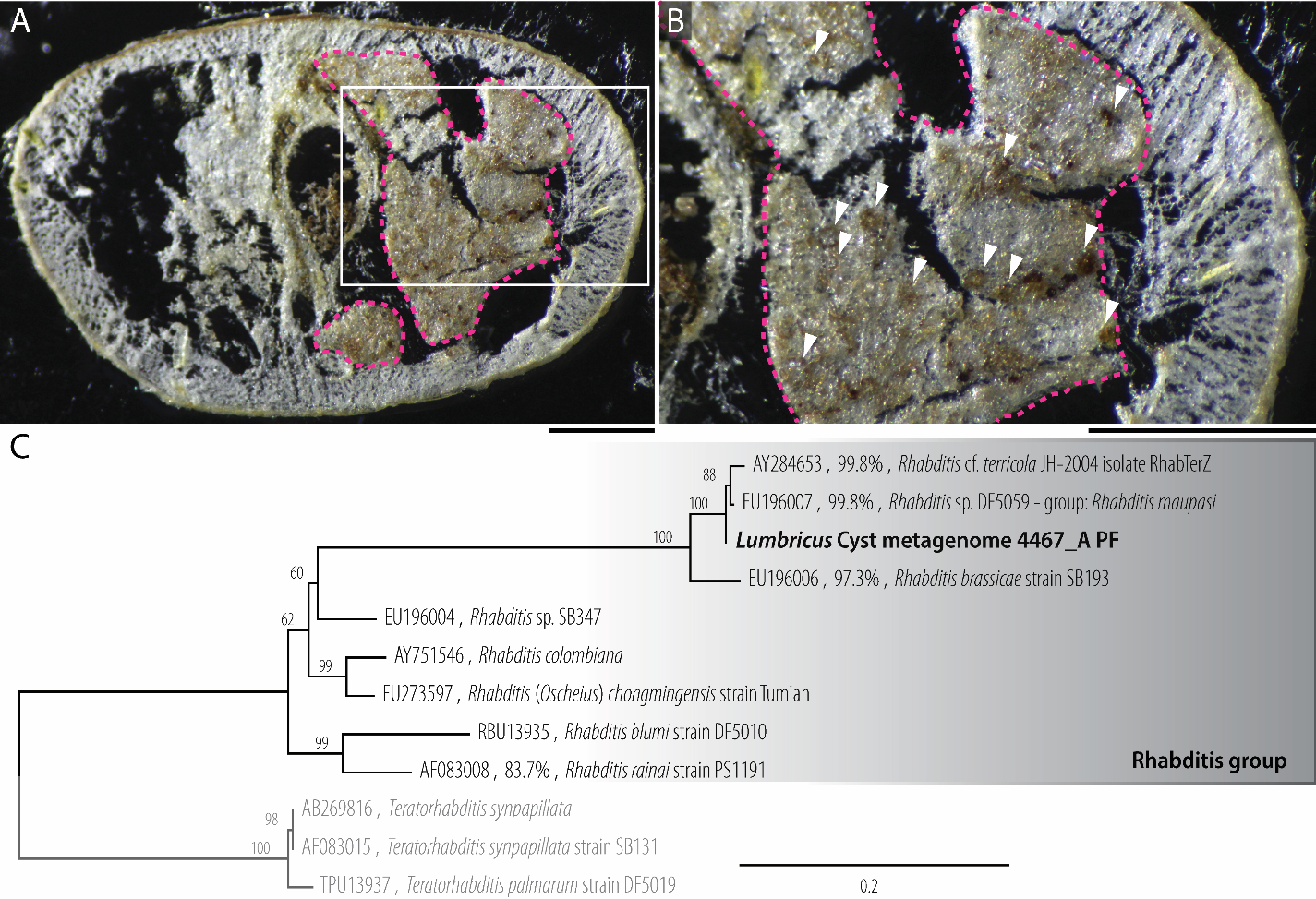

**Figure S-5: Tissue extraction of the nematode-containing brown bodies and phylogenetic analysis of the *Rhabditis* 18S rRNA gene from metagenomic sequencing. A)** Tissue of the brown bodies that encysted the nematodes (magenta outline) that was scraped off the glass slide for DNA extraction and metagenomic sequencing. **B)** Magnification (white box in A) shows cross-sections of the nematodes (consecutive section of the tissue section s3), indicated with white arrows. Microscopy scale bars: 300 µm **C)** The analysis shows that the nematodes were *Rhabditis maupasi*, known parasites that reside in the earthworms’ nephridia and coelomic cavity (e.g. *Lumbricus terrestris*, *Lumbricus rubellus*, *Allobophora longa*, and *Allobophora turgid*) (1). Numbers above the nodes indicate aLRT support values as estimated by fasttree2. Scale bar for tree shows substitutions per site.

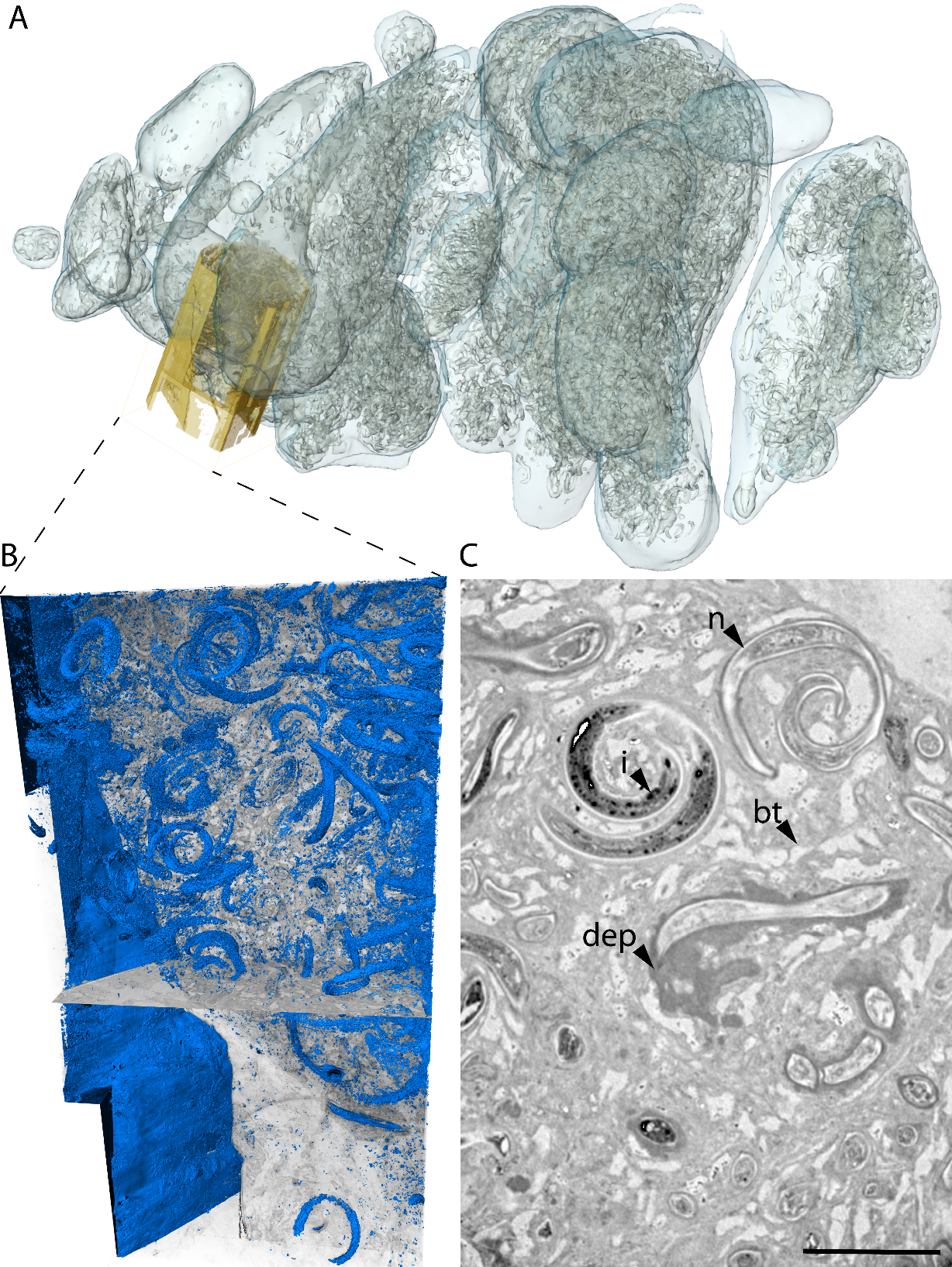

**Figure S-6: Microtomography of the nematodes. A)** Surface rendering of the nematode mass (grey outlines) within the brown bodies (transparent pale blue), with the co-registered higher resolution SRmicroCT dataset (yellow). **B)** Isosurface rendering of the magnified SRmicroCT nematode dataset, showing nematodes in blue and virtual slices through the *xy*- and *xz*-axes of the tissue stack. **C)** Magnified view of a virtual sectioning plane through the SRmicroCT data, showing different physiological states of the nematodes. **n**, nematode; **dep**, deposit; **bt**, brown body tissue; **i**, electron dense inclusions Scale bar 100 µm.

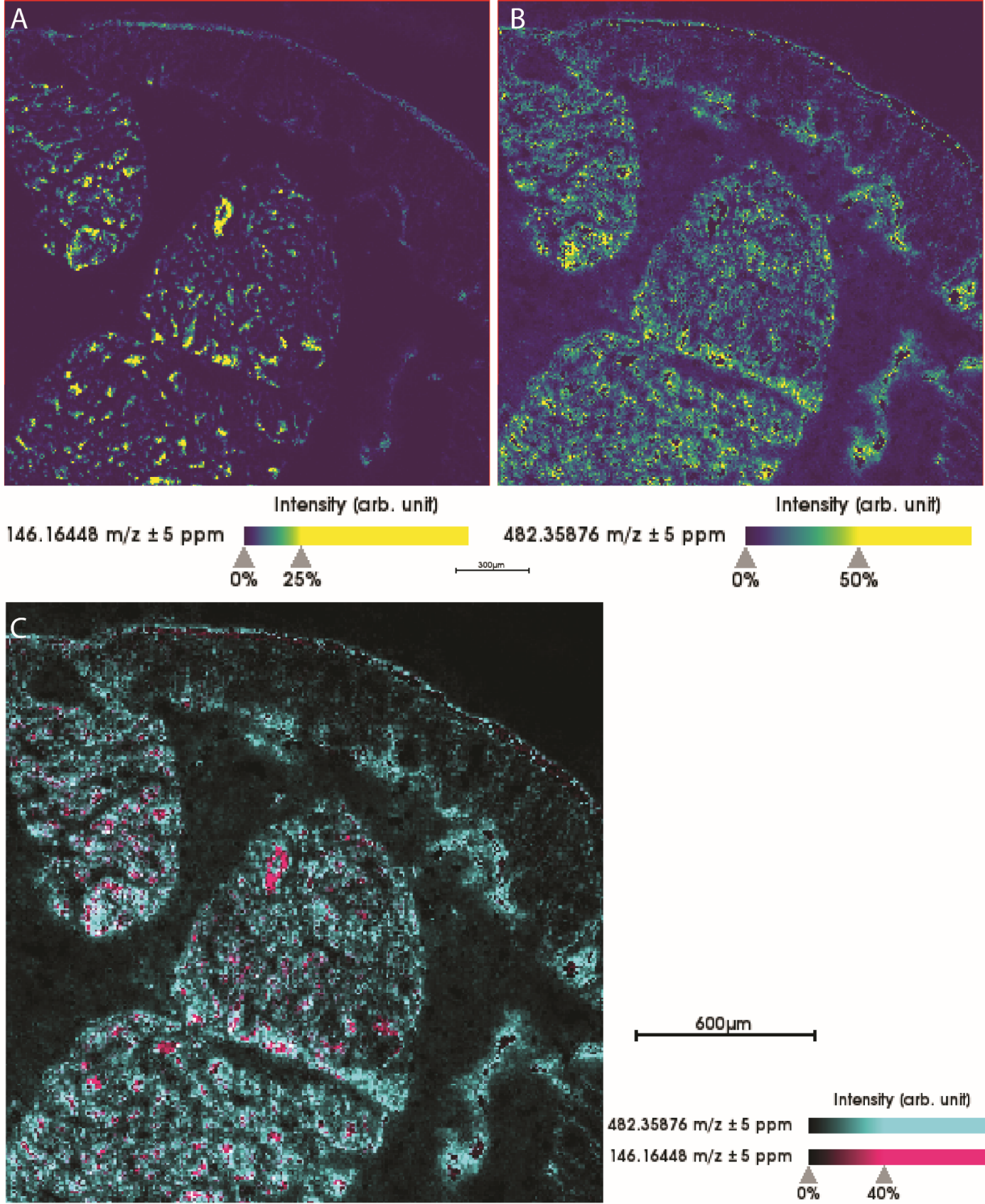

**Figure S-7: Ion maps showing the distributions of A) spermidine (*m/z* 146.1645) and B) PAF (*m/z* 482.3588), and C) an overlay of spermidine (magenta) and PAF (cyan)**. Full dataset of the magnified regions in Fig. 3, which show the distributions of the two compounds around nematodes.

**^++++^**
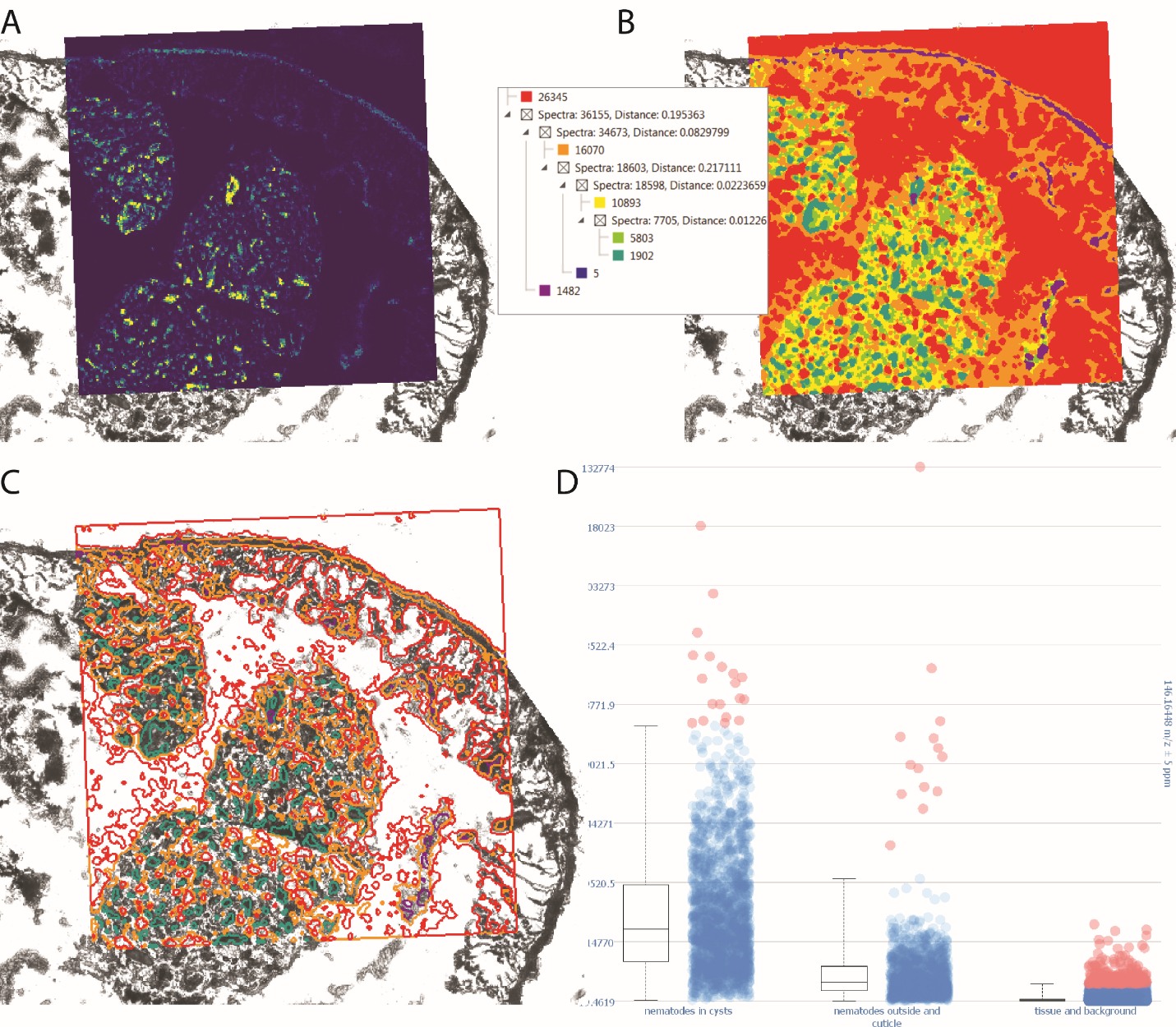

**Figure S-8: Quantification of the relative abundance of spermidine in the nematodes and the surrounding tissues**. **A)** Ion map of spermidine, shown in the ‘viridis’ color scheme. **B)** Spatial clustering of the high-resolution AP-MALDI-orbitrap-MSI dataset using a ‘weak’ smoothing option to reduce noise of the spatial clusters: red cluster, background; orange cluster, earthworm tissue (musculature); yellow and bright green clusters, tissue of the brown bodies (mainly hemocytes); dark green cluster, nematodes in cysts; blue and purple clusters, nematodes outside cysts and cuticle of the earthworm. One small region of a nematode within a cyst was assigned to a purple cluster (~10 pixel). **C)** Clusters were then converted into segmentation maps **D)** and used to quantify the relative abundance of spermidine in the nematode cysts (dark green cluster), outside the nematode cysts (purple and blue clusters) and in the musculature and background (red and orange clusters).

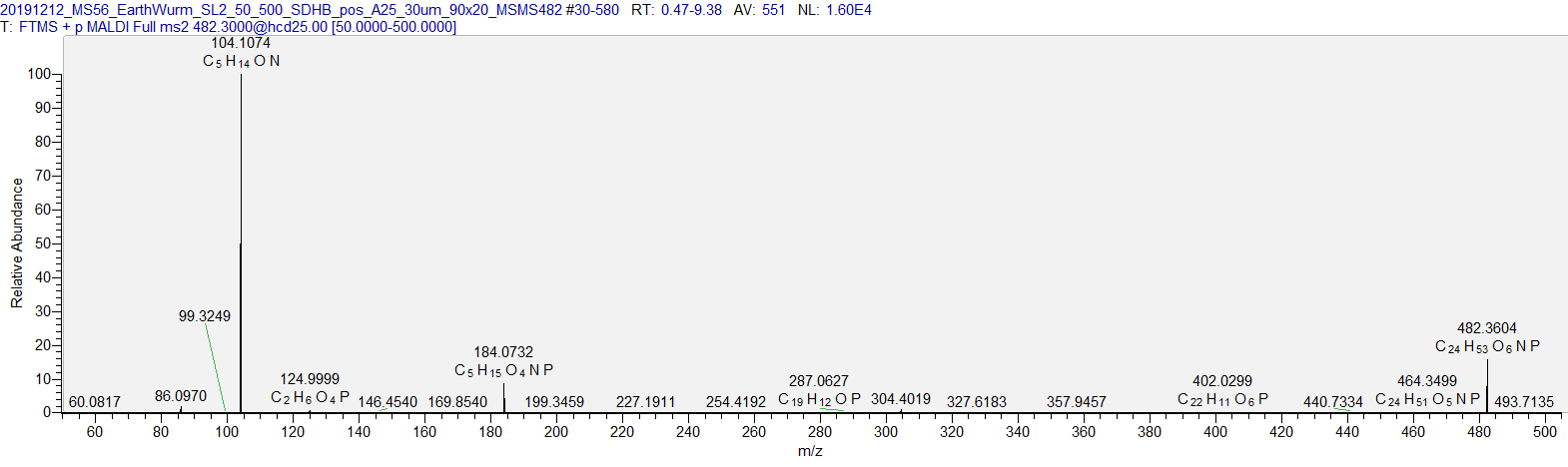

positive mode, HCD 25 constantly measured over a 90x20 area, raster width 30 µm

mass tolerance within 5ppm

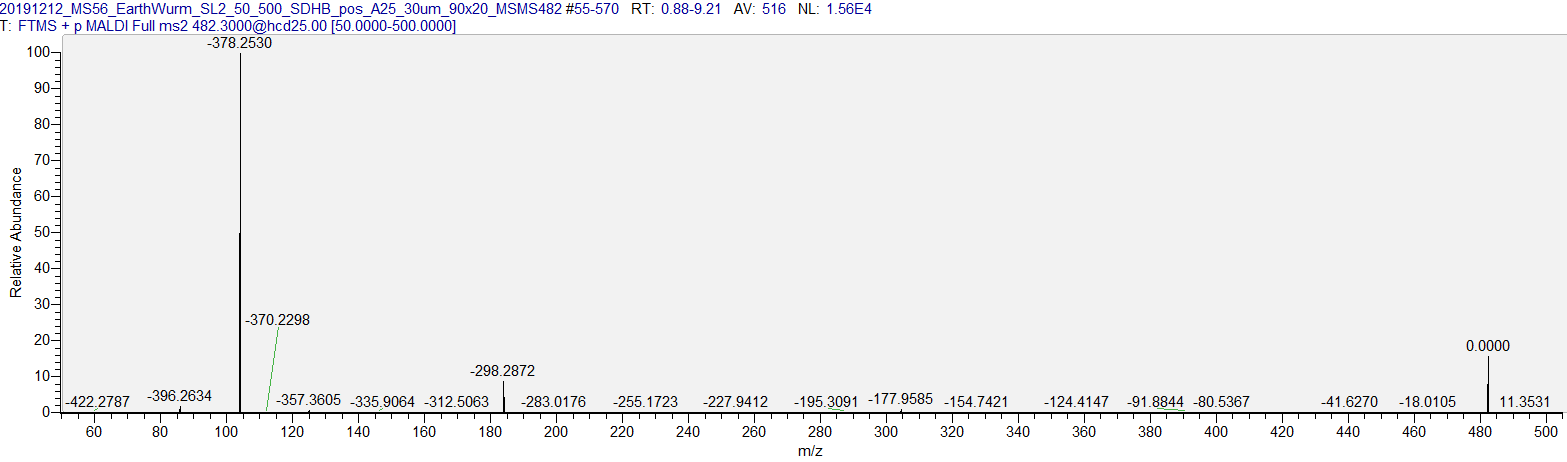

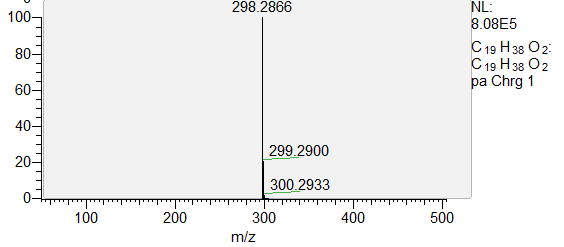

**Figure S-9:** **On-tissue** **MS^2^ spectra of of PAF (*m/z* 482.3605)**

We did on-tissue MS^2^ to verify METASPACE annotations of lysophosphatidylcholine O-16:0 ([C_24_H_52_NO_6_P + H]⁺, FDR 20%). Measurements were carried out in positive mode, with an HCD collision energy of 25 constantly measured over an area of 90 × 20 pixels with a raster width of 30 µm. The tolerance of the mass window was 5 ppm. For fragment ions, see *SI Appendix*, Table S-1 and methods for fragmentation procedures.

|  |  |  | Precursor ion | | | | | |  |  |  |  |  |  |  |
| --- | --- | --- | --- | --- | --- | --- | --- | --- | --- | --- | --- | --- | --- | --- | --- |
| Identification on METASPACE  [MPIMM_017_QE_P_LT](https://metaspace2020.eu/annotations?db=ChEBI-2018-01&grp=5727e853-e1dd-11e8-9d75-1b140a4756fa&ds=2019-09-25_17h41m59s) | **FDR** | **Detected *m/z* MALDI-MSI** | | **Calculated *m/z*** | **∆ ppm** | **Ion detected** | | **Formula** | **Detected *m/z* MALDI-MS/MS** | | **Calculated *m/z*** | **Formula fragment ion** | | **Charge state** | |
| Protoporphyrin, protoporphyrinate, protoporphyrin(2-) | 20% | 563.2661 | | 563.2653 | 0.8 | [M+H]+ | C_34_H_34_N_4_O_4_ | | - | - | | | - | | - |
| Lombricine, L-lombricine, D-lombricine | 20% | 271.0823 | | 271.0802 | 2.1 | [M+H]+ | | C_34_H_34_N_4_O_4_ | - | | - | - | | - | |
| Spermidine, spermidine(3+) | 20% | 146.1645 | | 146.1652 | 0.7 | [M+H]+ | | C_7_H_19_N_3_ | - | | - | - | | - | |
| lysophosphatidylcholine O-16:0/0:0,  2-hexadecyl-sn-glycero-3-phosphocholine (PAF) | 20% | 482.3604 | | 482.3605 | 0.2 | [M+H]+ | | C_24_H_52_NO_6_P | 184.0732  104.1074 | | 184.0733  104.1070 | C_5_H_14_O_4_NP  C_5_H_13_ON | | 1+  1+ | |
| Measurement of a metabolite standard |  |  | |  |  |  | |  |  | |  |  | |  | |
| Spermidine | - | 146.1652 | | 146.1652 | 0.0 | [M+H]+ | | C_7_H_19_N_3_ | 129.1387  112.1123 | | 129.1386  112.1121 | C_7_H_16_N_2_  C_7_H_13_N | | 1+  1+ | |

**Table S-1. Identifications of metabolites, detected with MALDI-MSI, based on MS^1^ annotations in METASPACE (****2****) and MS^2^ on the tissue and a standard (see methods).**
